# Pathological TDP-43 filaments accumulate at synapses and cause synaptic dysfunction

**DOI:** 10.64898/2026.01.27.701787

**Authors:** Renren Chen, Imogen Stockwell, Jessica C. Pierce, Sew-Yeu Peak-Chew, Melissa Huang, Kathy Newell, Bernardino Ghetti, Michael A. Cousin, Ingo H. Greger, Benjamin Ryskeldi-Falcon

## Abstract

The assembly of TAR DNA-binding protein 43 (TDP-43) into amyloid filaments within neurons is a hallmark of multiple neurodegenerative diseases, including motor neuron diseases (MND), frontotemporal dementias (FTD) and limbic-predominant age-related TDP-43 encephalopathy (LATE). These diseases result from the deterioration and loss of neurons, with synaptic dysfunction and neuronal hyperexcitability being prominent early events. Pathogenic mutations in the TDP-43 gene, *TARDBP*, that promote filament formation have established a causal role for TDP-43 assembly in neurodegenerative diseases. However, the molecular mechanisms underlying filament accumulation and their contribution to neurodegeneration are poorly understood. TDP-43 filaments can propagate between neurons in a prion-like manner, which may underlie the progressive spread and accumulation of TDP-43 pathology in disease. Here, we studied early stages of TDP-43 filament accumulation following internalisation of patient-derived TDP-43 filaments by mouse and human cortical neurons. Using proximity labelling, we identified molecular environments and putative interactions of TDP-43 filaments. We found that TDP-43 filaments accumulated at synapses, particularly in proximity to the presynaptic active zone, which we confirmed in FTD patient brain sections. Electron cryo-tomography (cryo-ET) directly visualised abundant TDP-43 filaments spanning the presynaptic cytoplasm *in situ*, which contacted synaptic vesicles and the plasma membrane. Functional measurements revealed that the accumulation of TDP-43 filaments led to presynaptic dysfunction and subsequent neuronal hyperexcitability. These findings suggest that synapses are a major early site of TDP-43 filament accumulation, relevant to their propagation, and directly link TDP-43 filament gain of function to synaptic dysfunction.

## INTRODUCTION

Assembly of the nucleocytoplasmic-shuttling, RNA-binding protein TDP-43 into filaments in neurons characterises multiple types of frontotemporal lobar degeneration (FTLD-TDP Types A-E); most types of MND, including amyotrophic lateral sclerosis (ALS); and the late-onset dementia LATE^1–3^. TDP-43 assembly is also a prevalent co-pathology in Alzheimer’s disease and Lewy body dementia^4^. Cryo-EM structures of TDP-43 filaments have established that they are amyloids, defined by the stacking of an ordered core protein fold stabilised by intermolecular β-sheets^5^. The core folds are composed of the N-terminal half of the TDP-43 low-complexity domain, with the remaining regions of TDP-43 flanking the core. Strikingly, while TDP-43 can adopt a wide range of filament core folds *in vitro*^6–8^, single distinct folds characterise different diseases^5,9,10^. TDP-43 filaments are specifically marked by aberrant post-translational modifications (PTMs), including truncation of the N-terminal flanking region and phosphorylation of the C-terminal flanking region at serine residues 409 and 410 (refs^1,2,11^). In postmortem brain, the filaments are concentrated within cytoplasmic inclusion bodies and, in certain diseases, within nuclei^12,13^. In addition, diffusely distributed TDP-43 filaments, commonly referred to as pre-inclusions, are present independently of inclusion bodies^14–17^. Pre-inclusions may represent early stages in filament accumulation, since they are less-well marked by ubiquitin and p62 than inclusions^13–15^. The subcellular localisation of pre-inclusions is poorly understood, although a subset in the neuropil colocalise with synaptic markers^18–22^.

Pathogenic mutations in the TDP-43 gene, *TARDBP*, that promote filament accumulation have established a causal role for this process in disease^23,24^. However, the molecular events that lead to filament accumulation are unknown. Postmortem staging has shown that TDP-43 filaments first accumulate in focal, disease-specific brain locations and subsequently progress to connected CNS regions^16,25,26^. This progressive accumulation mirrors patterns of synapse and neuron loss, and clinical symptoms^27–29^. Mounting evidence from cell and mouse models suggests that the prion-like propagation of TDP-43 filaments among neurons may account for their progressive accumulation^30–32^. This encompasses neuronal uptake, intracellular trafficking and release of TDP-43 filaments, in addition to amplification of filaments by seeded assembly of native TDP-43. However, the molecular environments and interactions that mediate TDP-43 filament propagation have not been investigated.

The mechanisms by which the accumulation of TDP-43 filaments results in neurodegeneration are also poorly understood. Synaptic dysfunction and neuronal hyperexcitability are early events in neurodegenerative diseases^33^, including TDP-43 proteinopathies, although how these relate to TDP-43 pathology is unclear. In postmortem brain, a proportion of neurons and glia with TDP-43 filament inclusions exhibit depletion of physiological nuclear TDP-43 (ref^1^), accompanied by loss of nuclear TDP-43 function effects, including mRNA mis-splicing^34^. Genetic risk variants in *UNC13A* that exacerbate its mis-splicing support a role for this effect in disease^35^. Induction of TDP-43 inclusions in cell and mouse models is sufficient to deplete nuclear TDP-43 and give rise to mis-splicing^36–44^, suggesting that TDP-43 filament accumulation drives loss of nuclear TDP-43 function in disease. In addition, multiple studies have reported toxic gain of function effects of TDP-43 inclusions in cell and mouse models^30,45–51^. However, the underlying mechanisms are unknown.

Here, to gain insight into TDP-43 filament accumulation, propagation and toxicity, we carried out proximity labelling of pathological TDP-43 filaments following their uptake by cultured neurons. This unbiased approach revealed molecular environments and putative interactions of TDP-43 filaments. We found that TDP-43 filaments accumulated at synapses and detected a pronounced enrichment of presynaptic active zone proteins, which we directly visualised *in situ* using cryo-ET. Assessment of synaptic vesicle dynamics, electrophysiological profiles and network-level calcium dynamics revealed that the accumulation of TDP-43 filaments led to synaptic dysfunction and subsequent network hyperexcitability. These findings suggest that synapses are a major early site of TDP-43 filament accumulation and directly link filaments to synaptic dysfunction and network hyperexcitability through a toxic gain of function.

## RESULTS

### Internalisation of patient-derived TDP-43 filaments by cultured neurons

To study early stages of TDP-43 filament propagation and accumulation within neurons, we took advantage of their ability to spontaneously internalise amyloid filaments (Figure 1A)^40,42,43,45,52,53^. This enabled us to load neurons with TDP-43 filaments in the absence of artificial protein delivery reagents and overexpression. We used TDP-43 filaments extracted from FTLD-TDP Type A patient brain (Supplementary Table 1), the most common FTLD, using the method we previously developed for cryo-EM structural studies (Methods)^5^. This ensured that we studied TDP-43 filaments with disease-relevant structures and PTMs^9^. Immunogold negative-stain electron microscopy confirmed that the extracted TDP-43 was filamentous (Figure 1B). Immunoblots confirmed that the filaments were composed of a mixture of full length and N-terminally truncated TDP-43 and were phosphorylated at serine residues 409 and 410 (pS409/410) (Figure 1C).

**Figure 1:**
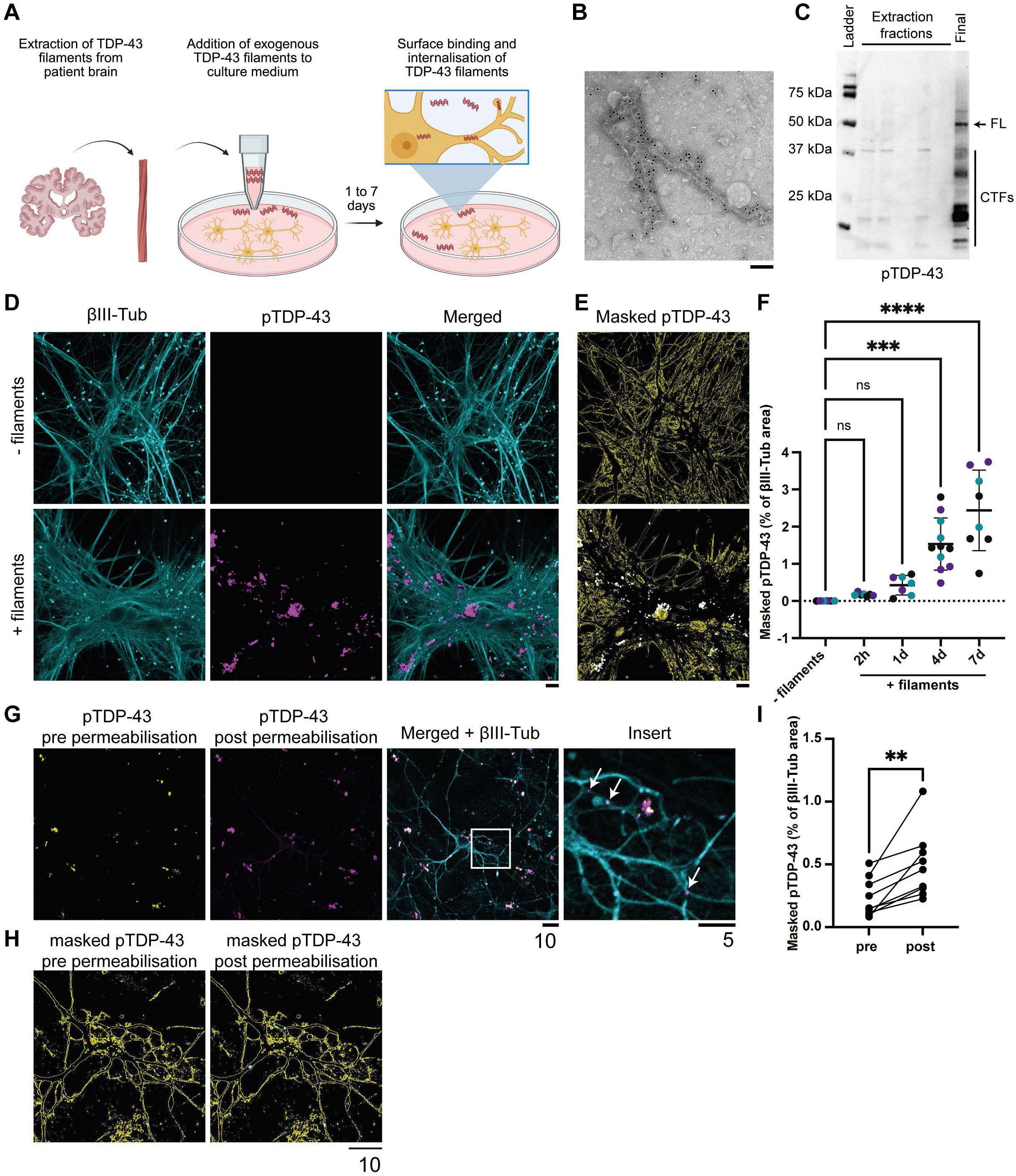
Internalisation of patient-derived pathological TDP-43 filaments by cultured neurons. **A.** Schematic summarising the extraction of TDP-43 filaments and their incubation with neuronal cultures. **B.** Immunogold negative-stain electron microscopy of extracted TDP-43 filaments from FTLD-TDP Type A patient brain, using a primary antibody against the N-terminus of TDP-43 and a secondary antibody conjugated to 10 nm gold particles. Scale bar, 100 nm. **C.** Immunoblot of extraction fractions and the final TDP-43 filament sample from FTLD-TDP Type A patient brain, probed with an antibody against pS409/410 TDP-43 (pTDP-43). The arrow indicates full-length (FL) TDP-43 and the bar indicates C-terminal fragments (CTFs) of TDP-43. **D.** Immunofluorescence confocal microscopy images of mouse primary cortical neurons incubated with (+ filaments) or without (- filaments) FTLD-TDP Type A patient-derived TDP-43 filaments for 7 d. Neurons were labelled using an antibody against class III β-tubulin (βIII-Tub; cyan) and TDP-43 filaments were labelled using the antibody against pTDP-43 (magenta). **E.** pTDP-43 signal (white) within images masked using the βIII-tubulin signal (yellow). Signal of 0.05-3 μm^2^ is shown in cyan. This size filter was used to exclude noise and large extracellular TDP-43 filament accumulations from quantification in **(F)**. Scale bar, 20 µm. **F.** Quantification of βIII-tubulin-masked pTDP-43 signal from **(E)** at different timepoints. The total area of pTDP-43 signal as a percentage of the βIII-tubulin signal mask is plotted. Each data point represents one technical repeat. Data points are colour coded by biological replicate. n = 3 biological replicates. Means +/- SD are shown. A one-way ANOVA with Tukey’s multiple comparison test was performed, ***p<0.001, ****p<0.0001. **G.** Immunofluorescence confocal microscopy images of mouse primary cortical neurons incubated with FTLD-TDP Type A patient brain-derived TDP-43 filaments for 1 d. TDP-43 filaments were labelled with the antibody against pTDP-43 pre- (yellow) and post- (magenta) detergent permeabilisation. Neurons were labelled with an antibody against class III β-tubulin (βIII-Tub; cyan). Arrows indicate examples of pTDP-43 signal unique to permeabilised neurons. **H.** pTDP-43 signal (white) within images masked using the βIII-tubulin signal (yellow). Signal of 0.05-3 μm^2^ shown in cyan. This size filter was used to exclude noise and large extracellular TDP-43 filament accumulations from quantification in **(I)**. Scale bars, 5 and 10 μm, as indicated. **I.** Quantification of βIII-tubulin-masked pTDP-43 signal pre- and post-permeabilisation from **(H)**. The total area of pTDP-43 signal as a percentage of the βIII-tubulin signal mask is plotted. Each pair of data points represent one technical repeat. n = 3 biological replicates. A paired t-test was performed, **p<0.01.

We added pathophysiological concentrations of TDP-43 filaments (estimated 160 pM) to the culture medium of primary cortical neurons from wild-type CD1 mice at 14 d *in vitro* (Supplementary Figure 1). The cultures expressed markers of mature neurons, including MAP2, NeuN and βIII-tubulin (Supplementary Figure 2A), and produced both spontaneous and evoked synaptic currents (Supplementary Figure 2B and C). Immunofluorescence (IF) confocal imaging using an antibody against pS409/410 TDP-43 showed a time-dependent association of TDP-43 filaments with the neurons over 7 d (Figure 1D-F). No IF signal was detected in the absence of exogenous TDP-43 filaments, confirming the specificity of pS409/410 for TDP-43 filaments.

To distinguish between surface-associated and internalised filaments, we repeated the IF prior to and following detergent permeabilisation of the neurons (Figure 1G-I). A proportion of TDP-43 filaments could be detected prior to permeabilisation, showing that they were associated with the surfaces of neurons. An additional population of TDP-43 filaments could only be detected following permeabilisation, confirming that they were internalised. Internalised filaments formed puncta in the cytoplasm of neuronal soma and neurites, reminiscent of pre-inclusions^14–17^.

We repeated these experiments in wild-type human cortical-like excitatory neurons differentiated for 11-14 d from the H9 embryonic stem cell (ESC) line using a doxycycline-inducible neurogenin-2 transcription factor cassette^54,55^. The cultures expressed markers of mature neurons, including βIII-tubulin (Supplementary Figure 2D). Similar to our results using mouse primary cortical neurons, IF confocal imaging showed the time-dependent association of TDP-43 filaments with the neurons (Supplementary Figure 2D-F). IF prior to and following detergent permeabilisation confirmed the presence of both surface-associated and internalised filaments, the latter of which formed cytoplasmic pre-inclusion-like puncta in neuronal soma and neurites (Supplementary Figure 2G and H).

The formation of TDP-43 filament inclusion bodies can induce the depletion of nuclear TDP-43 (ref^36^). To test for depletion of endogenous nuclear TDP-43 in the mouse primary neurons following incubation with exogenous filaments, we performed IF using an antibody against mouse TDP-43 and quantified the fluorescence intensity of nuclei (Supplementary Figure 3A and B). We found no significant difference between untreated cultures and cultures incubated with TDP-43 filaments for 7 d, suggesting no overt nuclear depletion at this timepoint. In conclusion, we produced two neuronal models of the cytoplasmic internalisation of brain-derived pathological TDP-43 filaments, in the absence of nuclear TDP-43 depletion.

### Antibody-targeted proximity labelling of TDP-43 filaments

To identify molecular environments and putative interactions of TDP-43 filaments in an unbiased manner, we used a time-resolved antibody-targeted proximity labelling approach (Figure 2A, Methods). After 1 and 3 d incubation with filaments, neurons were fixed, permeabilised and incubated with the primary antibody against pS409/410 TDP-43 to target proximity labelling to pathological filaments. Horseradish peroxidase conjugated to a secondary antibody was then used to catalyse the conversion of biotin-phenol to biotin-phenoxyl radicals in the presence of hydrogen peroxide, thereby biotinylating proximal biomolecules.

**Figure 2:**
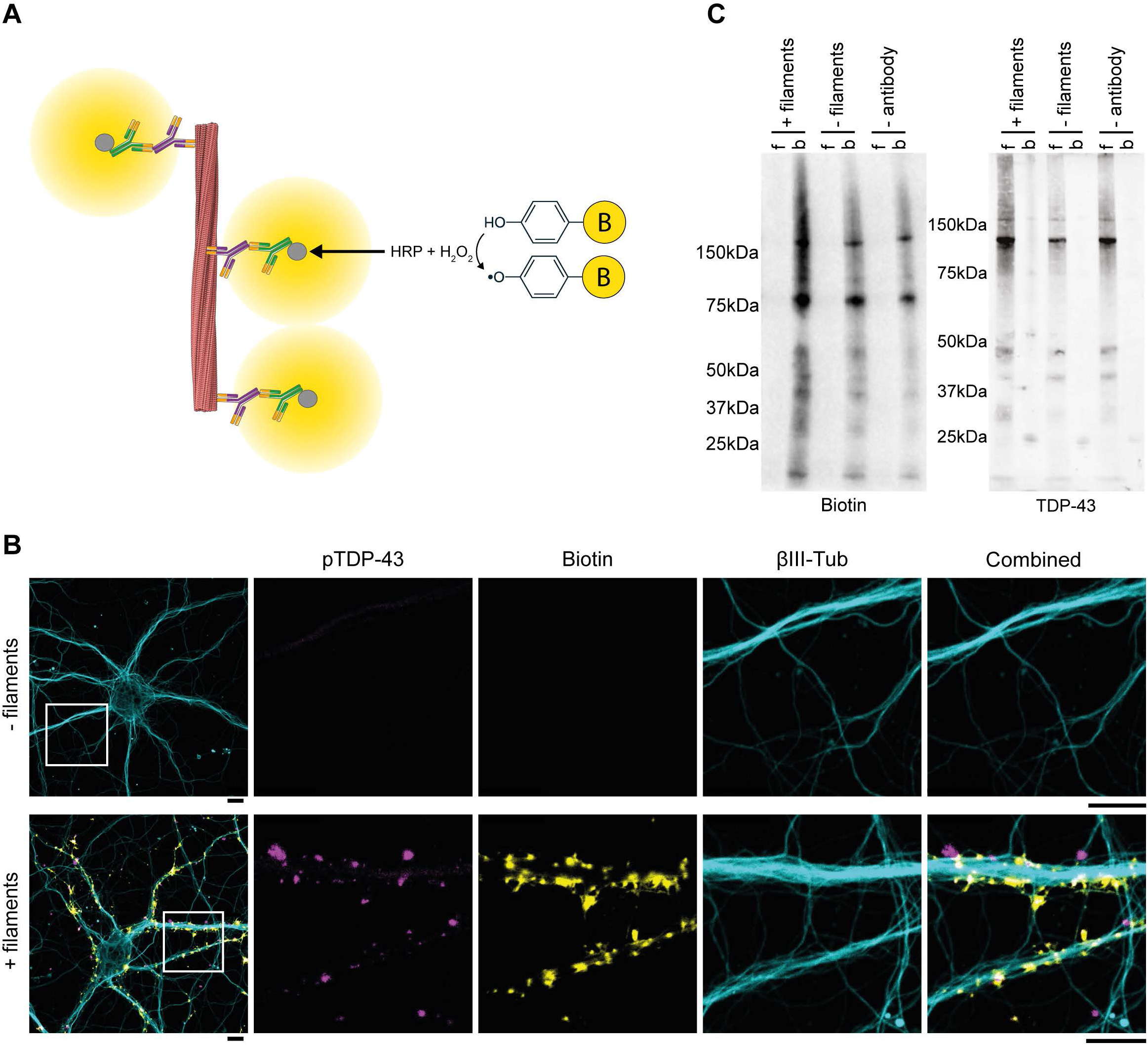
Antibody-targeted proximity labelling of TDP-43 filaments in cultured neurons. **A.** Schematic summarising the antibody-targeted proximity labelling strategy. TDP-43 filaments (red) were targeted using a primary antibody against pS409/410 TDP-43 (magenta). Horseradish peroxidase (grey) conjugated to a secondary antibody (green) was then used to catalyse the conversion of biotin-phenol to biotin-phenoxyl radicals (yellow) in the presence of hydrogen peroxide, thereby biotinylating proximal biomolecules. **B.** Immunofluorescence confocal microscopy images of mouse primary cortical neurons incubated with (+ filaments) or without (- filaments) FTLD-TDP Type A patient brain-derived TDP-43 filaments for 1 d, followed by antibody-targeted proximity labelling. TDP-43 filaments were labelled using an antibody against pS409/410 TDP-43 (pTDP-43; magenta), biotin was labelled using fluorescence-conjugated streptavidin (yellow) and neurons were labelled with an antibody against class III β-tubulin (βIII-Tub; cyan). Scale bars, 10 μm. **C.** Immunoblots of affinity purified biotinylated proteins from mouse primary cortical neurons (beads, b) and the flowthrough (f), probed with antibodies against biotin (left) and TDP-43 (right). Neurons were incubated with (+ filaments) or without (- filaments) FTLD-TDP Type A patient brain-derived TDP-43 filaments, in the presence or absence (- antibody) of the antibody against pTDP-43, followed by antibody-targeted proximity labelling.

IF confocal imaging showed the successful deposition of biotin proximal to TDP-43 filaments in the mouse primary and human ESC-derived cortical neurons (Figure 2B and Supplementary Figure 4A). No biotin signal was detected in the absence of filaments, confirming antibody-targeting of proximity labelling specifically to TDP-43 filaments. Biotin signal was not observed in the nucleus, further confirming that physiological nuclear TDP-43 was not targeted. Immunoblots of the biotinylated proteins following their affinity purification using streptavidin showed an increased number of proteins compared to conditions in the absence of either filaments or primary antibody (Figure 2C and Supplementary Figure 4B). Analysis of the flowthrough showed that the purification was exhaustive. Immunoblots using an antibody against total TDP-43 showed that the majority of TDP-43 was not biotinylated and remained in the flowthrough (Figure 2C and Supplementary Figure 4B), further confirming the specificity of proximity labelling for filaments. This provides a method for selectively targeting proximity labelling to pathological TDP-43 filaments.

### Analysis of TDP-43 filament proximity labelling

We performed label-free mass spectrometry to identify and quantify the biotinylated proximal proteins purified from the mouse and human neurons after 1 and 3 d incubation with TDP-43 filaments (Methods and Supplementary Information 1-8). Experiments performed in the absence of either filaments or primary antibody were included in the analysis to control for possible contaminants. We also performed proximity labelling on TDP-43 filaments deposited on culture plates in the absence of neurons, to control for proteins that co-extracted with the filaments from human brain (Supplementary Figure 4C). For both mouse primary and human ESC-derived neurons, the number of significantly enriched (*p*<0.05) proximal proteins increased between 1 and 3 d (90 to 164 for mouse and 154 to 470 for human), with <10% overlap between the timepoints (10 for mouse and 61 for human) (Figure 3A, Supplementary Data). Surprisingly, there was only a small overlap between human and mouse cultures at both timepoints (21 at 1 d and 45 at 3 d).

**Figure 3:**
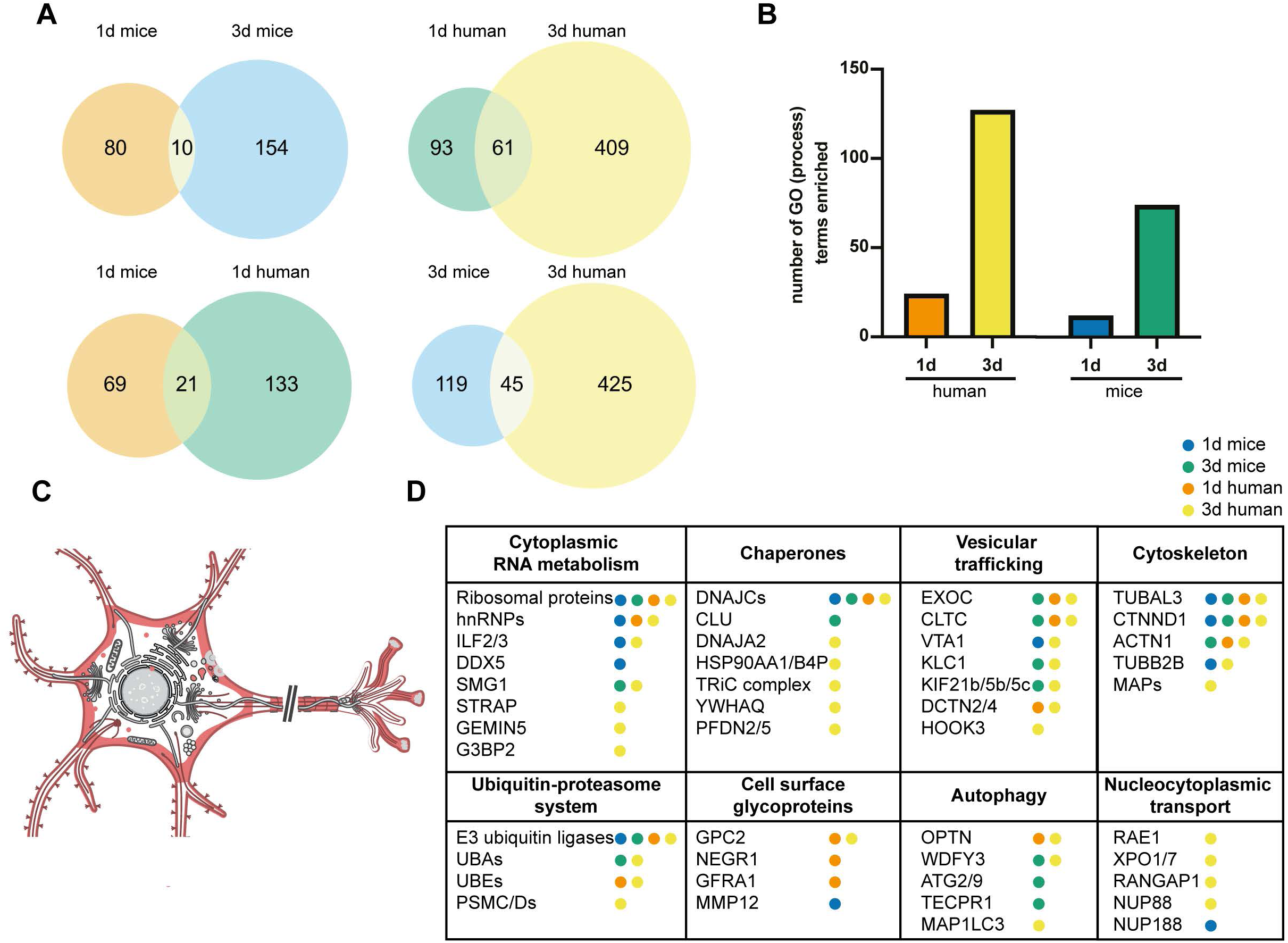
Analysis of antibody-targeted proximity labelling of TDP-43 filaments in cultured neurons. **A.** Venn diagrams of the number of significantly-enriched proximal proteins from mouse primary cortical neurons and human ESC-derived cortical neurons incubated with FTLD-TDP Type A patient-brain derived TDP-43 filaments for 1 d and 3 d. **B.** Bar plot of numbers of significantly-enriched Gene Ontology (GO) Biological Process terms for proximal proteins from mouse primary cortical neurons and human ESC-derived cortical neurons incubated with FTLD-TDP Type A patient-brain derived TDP-43 filaments for 1 d and 3 d. **C.** Schematic summarising the significantly-enriched GO Cellular Compartment terms (red) for mouse primary cortical neurons and human ESC-derived cortical neurons incubated with FTLD-TDP Type A patient-brain derived TDP-43 filaments for 1 d and 3 d. **D.** Representative significantly-enriched proximal proteins from mouse primary cortical neurons and human ESC-derived cortical neurons incubated with FTLD-TDP Type A patient-brain derived TDP-43 filaments for 1 d and 3 d.

The increase in proximal proteins between 1 and 3 d is consistent with the time-dependent association of TDP-43 filaments with neurons (Figure 1 and Supplementary Figure 2), as well as with a dynamic process in which the filaments encounter additional molecular environments over time as they propagate. In support of the latter, Gene Ontology (GO) analysis showed an increase in the number of significantly enriched Biological Process terms between 1 and 3 d for both neuronal cultures (Figure 3B and Supplementary Data). Consistent with our IF imaging (Figure 1 and Supplementary Figure 2), analysis of significantly enriched GO Cellular Component terms showed that the proximal proteins were associated with the cell surface, cytoplasm, axons, dendrites and synapses at both timepoints for both neuronal cultures (Figure 3C and Supplementary Data). GO terms relating to the nucleus were not significantly enriched, further supporting the specificity of proximity labelling for cytoplasmic TDP-43 filaments over physiological nuclear TDP-43.

Combining the GO analysis with manual inspection of the significantly enriched proximal proteins indicated that the filaments were associated with nine broad molecular environments: Cytoplasmic RNA metabolism, the cytoskeleton, nucleocytoplasmic transport, vesicular trafficking (including endocytosis, transport and exocytosis), molecular chaperones, the ubiquitin proteasome system, autophagy and cell-surface glycoproteins (Figure 3D and Supplementary Data). Cell surface glycoproteins and proteins involved in vesicular trafficking were enriched at both timepoints for both cultures. Proteins associated with the ubiquitin proteasome system and chaperones were less enriched at 1 d compared to 3 d for both cultures. Autophagy-related proteins were enriched at both timepoints in human neurons but were only enriched at 3 d for mouse neurons. Proteins associated with cytoplasmic RNA metabolism were enriched at 1 d in mouse neurons, but 3 d in human neurons. IF for proteins representative of these processes in the human ESC-derived cortical neurons after 3 d incubation with filaments confirmed colocalisation with TDP-43 filaments, validating the proximity labelling (Supplementary Figure 5). This reveals that TDP-43 filaments undergo dynamic localisation to multiple molecular environments following their binding to and internalisation by cultured neurons.

### Synaptic accumulation of TDP-43 filaments

There was a striking significant enrichment of synaptic proximal proteins, comprising both pre-and postsynaptic proteins, for both the mouse and human neurons at 1 and 3 d (Figure 4A and Supplementary Data). In agreement, synaptic terms were the most enriched across the GO analysis (Supplementary Data). This was especially pronounced for presynaptic active zone terms, with a 32-fold enrichment of the Cellular Component term ’Presynaptic Active Zone’ in the GO enrichment analysis of the mouse primary neurons at 3 d (Supplementary Data). All of the core components of the presynaptic active zone complex, comprising liprin-α, rab3-interacting molecule (RIM), RIM-binding protein (RIM-BP), uncoordinated 13 (unc13), ELKS-rich protein (ELKS), piccolo, bassoon and calcium/calmodulin-dependent serine protein kinase (CASK), were significantly enriched in the mouse and human neurons proximity labelling, with the sole exception of RIM-BP (Figure 4A)^56^. Synaptic vesicle-associated proteins were also abundant, including synaptic vesicle glycoprotein 2A, synaptic vesicle transporters (vesicular GABA transporter and neurotransmitter transporter 4), the rabconnectin-3 complex, endophilin A2 and β-synuclein (Figure 4A). These results show that TDP-43 filaments overwhelmingly accumulated at synapses following their neuronal internalisation, with a pronounced association with the presynaptic active zone.

**Figure 4:**
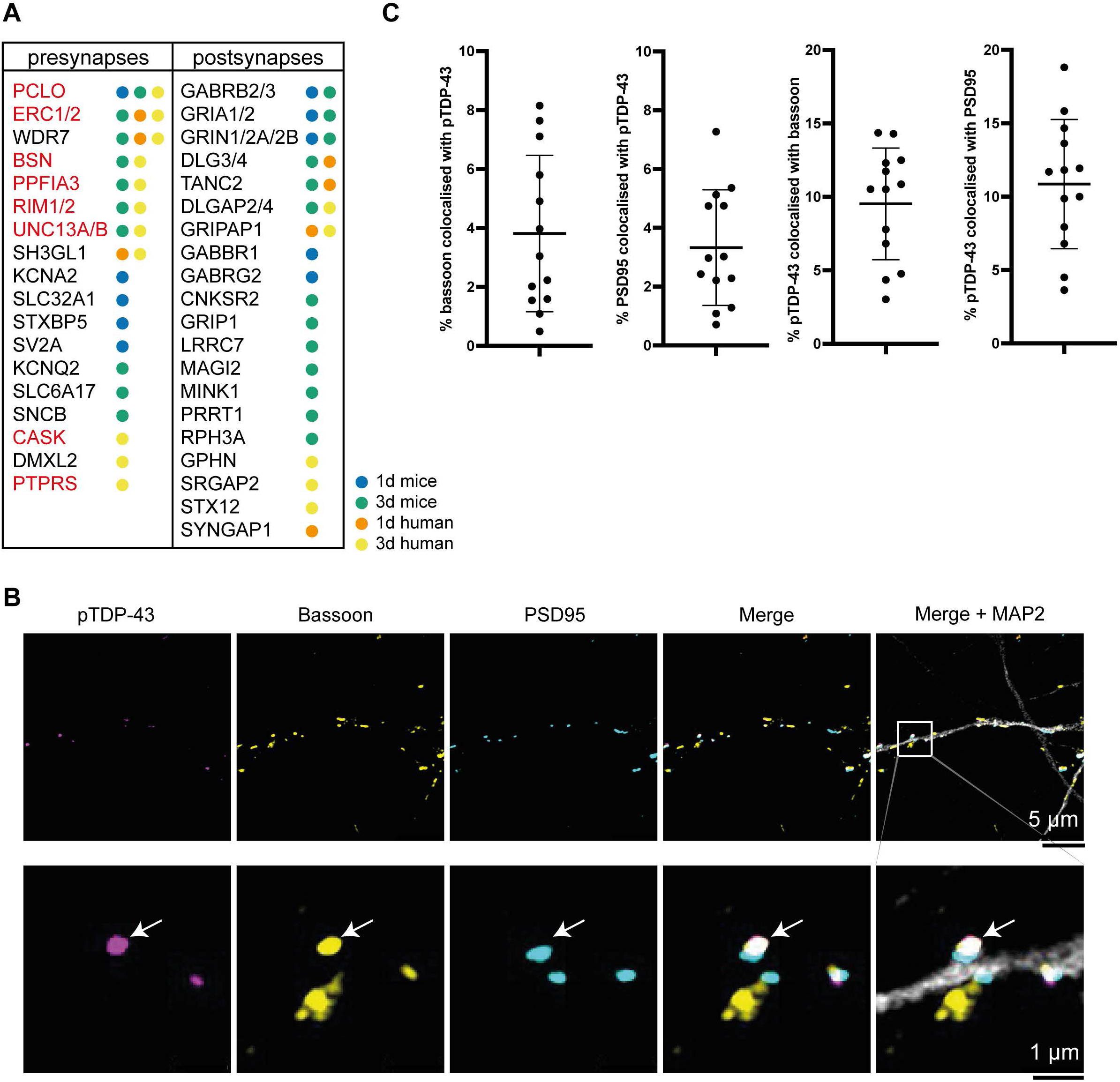
Synaptic accumulation of TDP-43 filaments in cultured neurons. **A.** Representative significantly-enriched proximal pre- and postsynaptic proteins from mouse primary cortical neurons and human ESC-derived cortical neurons incubated with FTLD-TDP Type A patient-brain derived TDP-43 filaments for 1 d and 3 d. Presynaptic active zone complex proteins are highlighted in red. **B.** Immunofluorescence Airyscan confocal microscopy images of mouse primary cortical neurons incubated with FTLD-TDP Type A patient-derived TDP-43 filaments for 3 d. Neurons were labelled using antibodies against microtubule-associated protein 2 (MAP2; grey); the presynaptic active zone complex protein bassoon (yellow); and PSD95 (cyan). TDP-43 filaments were labelled using the antibody against pTDP-43 (magenta). The arrow indicates an example of pTDP-43 signal colocalising with bassoon. Scale bars, 5 µm and 1 µm, as indicated. **C.** Percentage of MAP2-masked pTDP-43 signal colocalised with bassoon (left) and PSD95 (right), and vice versa, from **(C)** in three-dimensional space. Each data point represents one technical replicate. n= 2 biological replicates. Means +/- SD are shown.

IF Airyscan confocal microscopy of the neurons for pS409/410 TDP-43, the presynaptic active zone complex protein bassoon and the postsynaptic density protein 95 (PSD95) allowed us to confirm the accumulation of TDP-43 filaments at both the presynaptic active zone and the postsynaptic density, respectively (Figure 4B). An approximately equal proportion of pS409/410 TDP-43 signal (10%) colocalised with bassoon and PSD95 signal, affecting approximately 4% of total pre- and postsynapses (Figure 4C).

Immunoblots of synaptosomes isolated from mouse primary neurons after 3 d incubation with filaments using the antibody against pS409/410 TDP-43 confirmed the association of TDP-43 filaments with synapses (Figure 5A). Full length and N-terminally truncated pS409/410 TDP-43 were detected in synaptosome fractions from neurons that had been incubated with TDP-43 filaments, but not in neurons in the absence of filaments. Immunoblots using an antibody against the N-terminal half of TDP-43, which detects both pathological filaments and physiological TDP-43, indicated that physiological TDP-43 was also associated with synapses, irrespective of incubation with TDP-43 filaments (Figure 5A). This is consistent with previous reports of physiological TDP-43 at synapses^57–59^.

**Figure 5:**
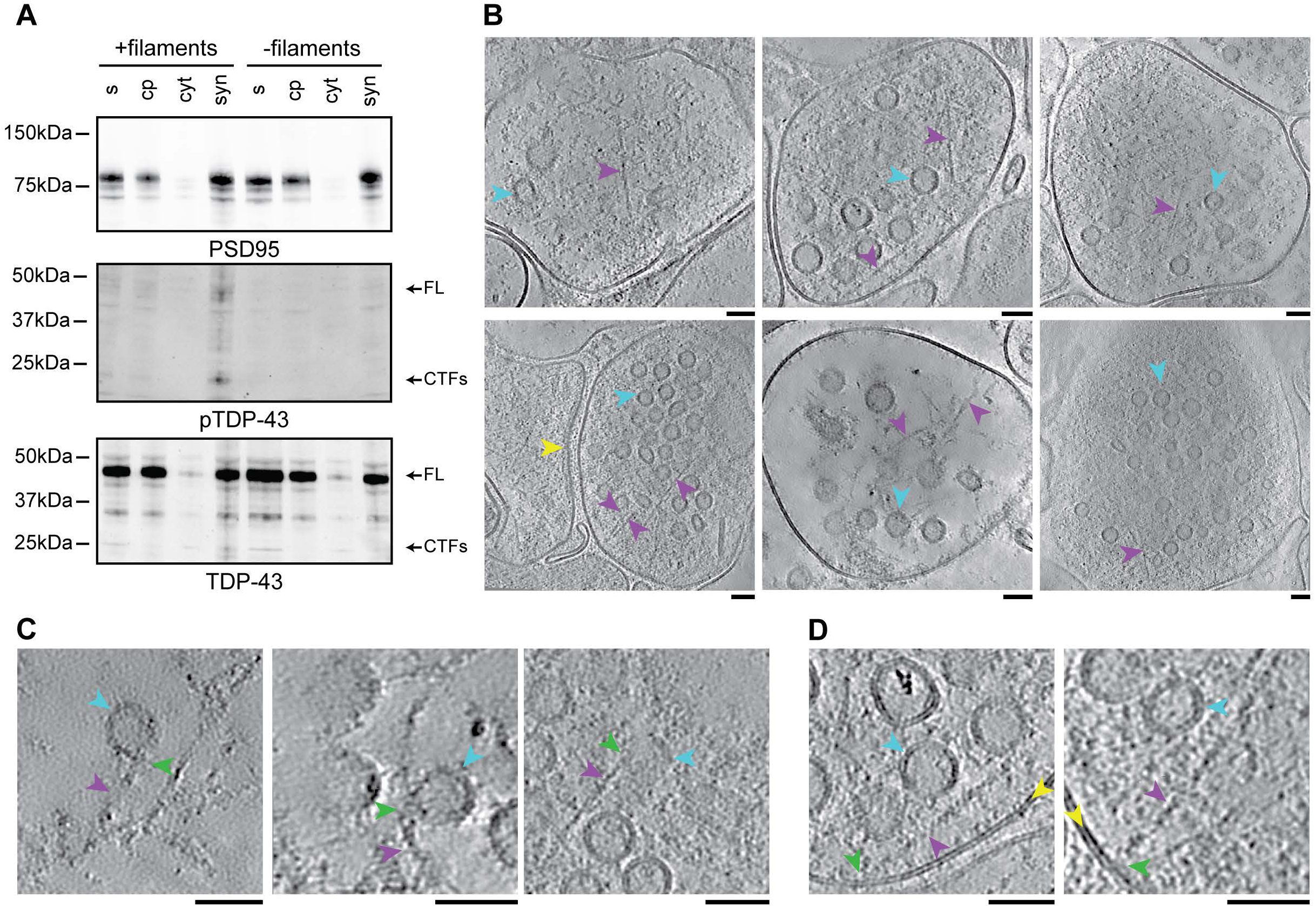
Cryo-ET of presynaptic TDP-43 filaments. **A.** Immunoblot of synaptosome isolation fractions (s, supernatant; cp, cell pellet; cyt, cytosolic; syn, synaptosomes) from mouse primary cortical neurons incubated with (+ filaments) and without (- filaments) FTLD-TDP Type A patient brain-derived TDP-43 filaments for 3 d, probed with antibodies against postsynaptic density protein 95 (PSD95, top), pS409/410 TDP-43 (pTDP-43, middle) and TDP-43 (bottom). The arrows indicate full-length (FL) TDP-43 and C-terminal fragments (CTFs) of TDP-43. **B.** Denoised tomographic slices of synaptosomes from mouse primary cortical neurons incubated with FTLD-TDP Type A patient brain-derived TDP-43 filaments for 3 d. TDP-43 filaments (magenta arrows), example synaptic vesicles (cyan arrows) and the postsynaptic density (yellow arrow) are indicated. **C and D.** Denoised tomographic slices showing TDP-43 filaments (magenta arrows). making contacts (green arrows) with synaptic vesicles (cyan arrows) **(C)** and the presynaptic plasma membrane (yellow arrows) **(D)**. **B-D.** Scale bars, 50 nm.

To further investigate the synaptic accumulation of TDP-43 filaments, we used cryo-ET to reconstruct molecular-resolution tomograms of the synaptosomes under near-native conditions (Figure 5B). This revealed abundant filaments in presynapses that matched the dimensions of the TDP-43 filaments^9^. These filaments were distinct from filamentous components of the presynaptic cytomatrix previously characterised using cryo-ET^60,61^, including F-actin, microtubules and synaptic vesicle connectors/tethers, which we also observed in addition to TDP-43 filaments (Supplementary Figure 6). The TDP-43 filaments spanned the presynaptic cytoplasm and were often in close proximity to synaptic vesicles and the plasma membrane. Closer inspection of these proximal sites revealed that some of the filaments made contact with synaptic vesicles and the plasma membrane (Figure 5C and D). We did not observe filaments contained within synaptic vesicles or other intercellular compartments such as endolysosomes. This direct visualisation of TDP-43 filaments within presynapses supports the proximity labelling, suggesting that TDP-43 filaments localise to and interact with components of synaptic vesicles and the presynaptic active zone.

To relate our findings to the human brain, we performed immunoblots for pS409/410 TDP-43 on the synaptosome fraction from FTLD-TDP Type A patient brain (Supplementary Figure 7A, Supplementary Table 1). As for the cultured neurons, full length and N-terminally truncated pS409/410 TDP-43 were detected in the synaptosome fraction. We also performed IF confocal microscopy and stimulated emission depletion (STED) super-resolution microscopy for bassoon and pS409/410 TDP-43 on FTLD-TDP Type A patient brain sections, which showed co-localisation in the neuropil (Supplementary Figure 7B–D). This is consistent with previous reports of co-localisation between TDP-43 pre-inclusions and pre- and post-synaptic proteins in FTLD-TDP and ALS^21,22^. Our neuronal models suggest that synaptic TDP-43 filament accumulation is an early stage in the propagation of TDP-43 pathology, with relevance to the human brain.

### Synaptic dysfunction

Given the synaptic accumulation of TDP-43 filaments and their proximity to the presynaptic active zone in our neuronal cultures, as well as in FTD and ALS patient brain^21,22^, we sought to understand if the filaments interfered with synaptic functions. We first determined the impact of TDP-43 filaments on synaptic vesicle recycling using the reporter synaptophysin-pHluorin (SypHy), a pH-sensitive enhanced green fluorescent protein (pHluorin) fused to the first intraluminal loop of the synaptic vesicle protein synaptophysin^62^. SypHy fluorescence is typically quenched in the acidic synaptic vesicle lumen and becomes unquenched upon synaptic vesicle fusion, resulting in an activity-dependent increase in fluorescence. Subsequently, SypHy is retrieved by synaptic vesicle endocytosis, with synaptic vesicle acidification re-quenching its fluorescence (Figure 6A).

**Figure 6:**
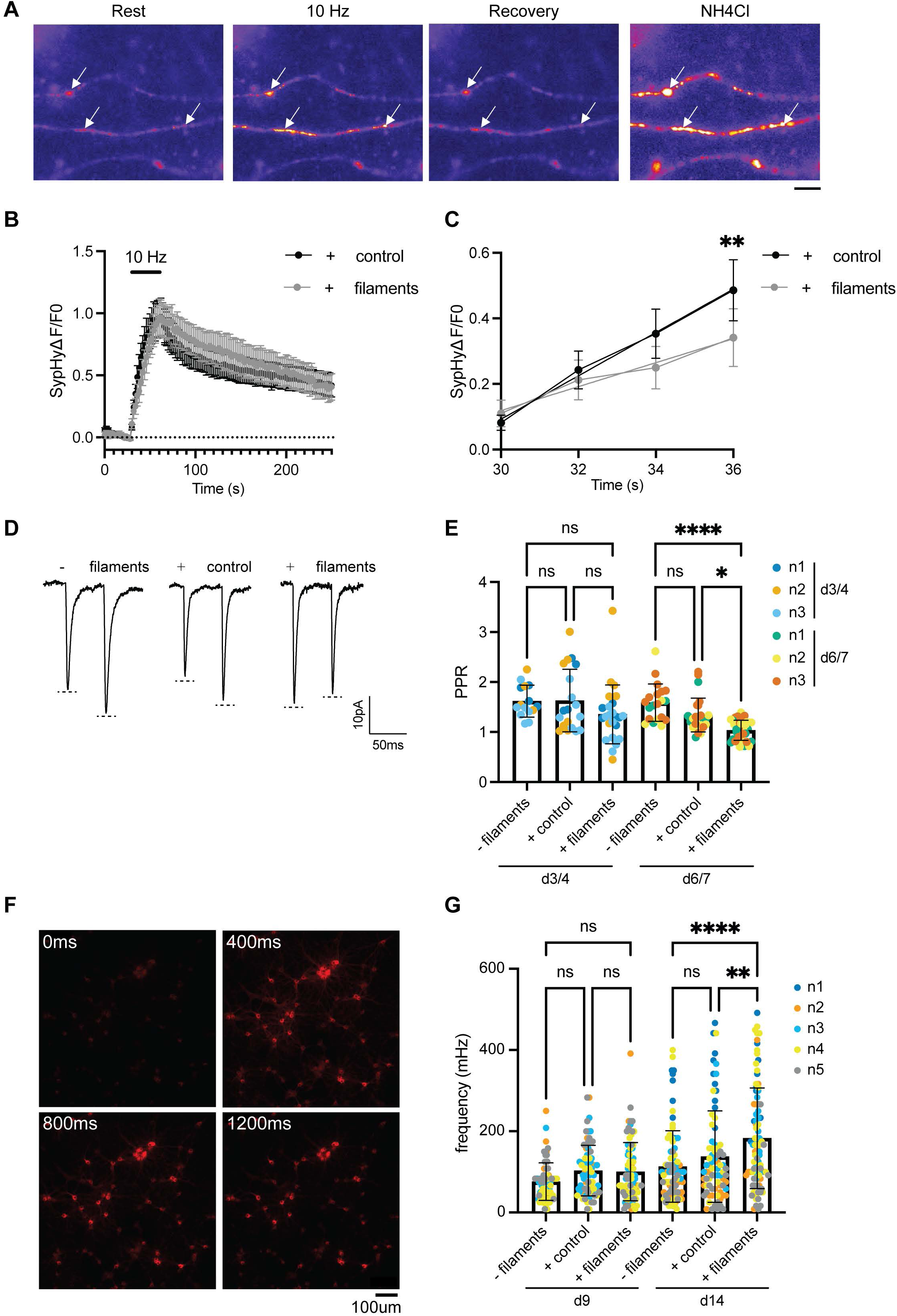
Presynaptic dysfunction and longer-term network hyperexcitability following TDP-43 filament accumulation in cultured neurons. **A.** Representative false-coloured images of mouse primary cortical neurons expressing SypHy prior to stimulation (Rest), during stimulation with a train of 300 action potentials delivered at 10 Hz (10 Hz), following stimulation (Recovery) and during challenge with ammonium chloride (NH_4_) buffer to reveal the total synaptic vesicle pool. Arrows indicate examples of terminals that respond to stimulation. Scale bar, 2 μm. **B.** Time course of relative SypHy fluorescence intensity (ΔF/F0) in mouse primary cortical neurons incubated with FTLD-TDP Type A patient brain-derived TDP-43 filaments (+ filaments) or with aged-matched control brain extracts (+ control) for 3 d, normalised to challenge with NH_4_ buffer (total synaptic vesicle pool). The bar indicates the period of stimulation. n = 4 biological replicates. Means +/- SEM are shown. **C.** Rate of activity-dependent relative SypHy fluorescence increase (ΔF/F0) over the first 6 seconds of stimulation (stimulation triggered at 30 s). n = 4 biological replicates. Means +/- SEM are shown. Linear regression analysis was performed (lines), **p=0.0055. **D.** Representative traces of excitatory postsynaptic currents (EPSCs) evoked by paired presynaptic stimuli (50 ms interval) from mouse primary hippocampal neurons incubated with (+ filaments) or without (- filaments) FTLD-TDP Type A patient brain-derived TDP-43 filaments or with aged-matched control brain extracts (+ control) for 7 d. **E.** Plot of the ratios of the second to the first EPSC (paired-pulse ratio; PPR) from mouse primary hippocampal neurons incubated with (+ filaments) or without (- filaments) FTLD-TDP Type A patient brain-derived TDP-43 filaments or aged-matched control brain extracts (+ control) for 3-4 d (d3/4) and 6-7 d (d6/7). Each data point represents one paired recording. Data points are colour coded by biological replicate. n = 3 biological replicates. Means +/- SD are shown. A two-way ANOVA with Tukey’s multiple comparison test was performed, *p<0.05, ****p<0.0001. **F.** Widefield fluorescence microscopy images of mouse primary cortical neurons expressing the fluorescent calcium indicator jRGECO1a (red) over 1200 ms. Images were taken at 100 ms intervals. Scale bar, 100 μm. **G.** The spontaneous firing frequency calculated from **(F)** for mouse primary cortical neurons incubated with (+ filaments) or without (- filaments) FTLD-TDP Type A patient brain-derived TDP-43 filaments or aged-matched control brain extracts (+ control) for 9 d (d9) and 14 d (d14). Each data point represents one time-lapse recording. Data points are colour coded by biological replicate. n = 5 biological replicates. Means +/- SD are shown. A two-way ANOVA with Tukey’s multiple comparison test was performed, **p<0.01, ****p<0.0001.

Mouse primary cortical neurons expressing SypHy were incubated for 3 d with either TDP-43 filaments or control extracts from aged human brains lacking amyloid filament pathology. SypHy fluorescence was then measured upon stimulation of the neurons using a train of 300 action potentials delivered at 10 Hz. The neurons displayed a characteristic response, with an evoked increase in fluorescence followed by a decrease post-stimulation (Figure 6B). However, there was a significant reduction in the initial rate of the fluorescence increase in neurons treated with TDP-43 filaments compared to those treated with control extracts (Figure 6C and Supplementary Table 1). Other parameters of the SypHy response, including peak fluorescence and fluorescence decay, were not significantly different between conditions (Supplementary Figure 8A and B). The number of responsive nerve terminals was also not affected by the presence of TDP-43 filaments (Supplementary Figure 8C). These results suggest that the coupling of synaptic vesicle fusion to neuronal activity was disrupted in neurons treated with TDP-43 filaments, without impacting the overall extent of synaptic vesicle fusion or endocytosis.

We next performed whole-cell electrophysiological recordings on synaptically connected pairs of mouse primary hippocampal neurons incubated with TDP-43 filaments. Hippocampal neurons were used instead of cortical neurons to facilitate detection of altered presynaptic properties, due to their homogeneous low release probabilities^63^. We used post-hoc IF Airyscan confocal microscopy to confirm that the mouse primary hippocampal neurons used in recordings accumulated TDP-43 filaments at presynapses (Supplementary Figure 9A), which occurred to a similar extend as we previously observed in mouse primary cortical neurons (Supplementary Figure 9B).

We performed paired-pulse recordings, in which we applied a pair of depolarising pulses, separated by 50 ms, to trigger action potentials in the presynaptic neuron that each evoked an excitatory current in the postsynaptic neuron (Figure 6D). We compared the ratio of the amplitude of the second response to that of the first, known as the paired-pulse ratio (PPR), among untreated neurons and neurons incubated with either TDP-43 filaments or control extracts (Figure 6E). The PPR relates to the probability of presynaptic neurotransmitter release and so can be used to assess presynaptic function^64^. We found a progressive decrease in the PPR (less paired-pulse facilitation) over 7 d in neurons incubated with TDP-43 filament extracts compared to untreated neurons and neurons incubated with control extracts. The amplitudes of the current response to the first stimulus were similar among all groups, suggesting that the decrease in PPR was not driven by differences in initial release quanta (Supplementary Figure 9C). The frequency and amplitude of spontaneous postsynaptic current responses were also similar among all groups (Supplementary Figure 9D). These results suggest that TDP-43 filaments reduce synaptic facilitation through presynaptic dysfunction, in agreement with the SypHy imaging (Figure 6C).

We also assessed spontaneous network-level calcium dynamics in mouse primary cortical neurons expressing the fluorescent calcium indicator jRGECO1a (Figure 6F, Supplementary Figure 10A). We observed a significant increase in the frequency of spontaneous synchronous calcium transients among neurons incubated with TDP-43 filaments for 14 d compared to untreated neurons and neurons incubated with control extracts (Figure 6G). This effect was not seen at earlier timepoints. The synchronous nature of this increase in calcium transient frequency suggests that it reflects a network hyperexcitability phenotype. The amplitudes and durations of the calcium transients were not significantly different between neurons treated with TDP-43 filaments and control brain extracts (Supplementary Figure 10B and C). These results suggest that TDP-43 filament accumulation, in the absence of nuclear depletion, leads to presynaptic dysfunction followed by network hyperexcitability in cultured neurons

## DISCUSSION

The assembly of TDP-43 into amyloid filaments is central to multiple neurodegenerative diseases. Prion-like propagation of TDP-43 filaments is thought to underlie their progressive accumulation in the brain. Evidence suggests that toxicity arises from sequestration of TDP-43 within filaments, leading to its loss of function, in addition to toxic gain of function of the filaments themselves. However, the molecular events underlying filament propagation and toxicity are poorly understood, which limits our ability to target them for possible therapeutic benefit.

We carried out antibody-targeted proximity labelling of TDP-43 filaments following their internalisation by cultured neurons, to investigate their molecular environments and putative interactions in an unbiased manner. We used TDP-43 filaments extracted from patient brain, because, to date, the sequences that form the ordered cores of TDP-43 filaments assembled *in vitro* and their folds have differed from those found in disease^6–8^. In addition, *in vitro*-assembled filaments lack disease-associated PTMs^1,2,11^. The use of an antibody against pS409/410 TDP-43 enabled us to target proximity labelling to native TDP-43 filaments. This provided a rich representation of the molecular environments of the filaments. As this approach identifies proximal proteins, putative interactions will need to be validated. Owing to the low abundance, large size and insolubility of TDP-43 filaments and other amyloids, proximity labelling may be better suited to studying their interactions than methods commonly used to study soluble proteins, such as co-immunoprecipitation.

Many of the significantly enriched proteins from our proximity labelling align with previously-reported putative interactions of cytoplasmic TDP-43 inclusions, including with proteins involved in cytoplasmic RNA metabolism^38,65–67^, nucleocytoplasmic transport^65–67^, the cytoskeleton^38,65–68^, vesicular trafficking^65,66^, molecular chaperones^38,65,66,68^, the ubiquitin proteasome system^65,66,69,70^ and autophagy^69–71^. Importantly, our study links these molecular environments to pathological TDP-43 filaments and their propagation. Proximity to cytoplasmic RNA metabolism factors, together with the fact that the TDP-43 RNA recognition motifs lie outside of the ordered filament core^9^, suggest that TDP-43 filaments may retain the ability to interact with these factors. This may be related to a recent report of the sequestration of additional splicing regulators by TDP-43 inclusions in a neuronal model, which resulted in mis-splicing^51^. Our identification of significantly enriched cell surface glycoproteins, and proteins involved in endocytosis, endolysosomal trafficking and exocytosis may be related to the binding, internalisation, intracellular transport and release of TDP-43 filaments by neurons, respectively, during propagation. Similar proteins have previously been linked to the propagation of neurodegenerative disease-associated filaments of tau and α-synuclein^72–74^. Thus, our proximity labelling results provide a rich dataset for future research, supporting previously reported interactions of pathological TDP-43, as well as suggesting several novel molecular environments and putative interactions with relevance to propagation.

Here, we focussed on the synaptic accumulation of TDP-43 filaments following their internalisation by neurons. Our proximity labelling revealed a striking number of significantly enriched synaptic proteins, most of which were components of the presynaptic active zone and synaptic vesicles^56^. Previous proximity and interaction studies of cytoplasmic TDP-43 inclusions did not detect synaptic proteins because they were conducted in non-neuronal cells. We confirmed the synaptic accumulation of TDP-43 filaments using IF imaging and synaptosome fractionation in the neuronal cultures and in FTLD-TDP Type A patient brain. Cryo-ET directly visualised TDP-43 filaments within presynapses under near-native conditions at molecular-resolution. The filaments spanned the presynaptic cytoplasm and often contacted synaptic vesicles and the plasma membrane. This provides a rationale for the enrichment of synaptic vesicle and presynaptic active zone proteins during proximity labelling. It is possible that some of these proteins identified by proximity labelling might mediate TDP-43 filament interactions with synaptic vesicles and the plasma membrane revealed by cryo-ET.

Our study links previous observations of synaptic TDP-43 pre-inclusions to the propagation of pathological TDP-43 filaments^21,22^. Synaptic transfer of filaments may account for the observation that filament accumulation is constrained within connected neuronal networks in disease^16,25,26^. We did not observe TDP-43 filaments within synaptic vesicles using cryo-ET, suggesting that these vesicles may not directly contribute to filament release and that alternative mechanisms may exist, perhaps involving extracellular vesicles^75^. The presence of physiological TDP-43 at synapses^57–59^, which we confirmed here, suggests that synapses may also represent a site of seeded assembly. While we studied initial stages of propagation following the internalisation of filaments by neurons, future work could use the models and approach presented here to gain insight into the molecular mechanisms of subsequent propagation stages.

We found that the internalisation and accumulation of TDP-43 filaments in primary neurons resulted in presynaptic dysfunction, which manifested as a reduced initial rate of synaptic vesicle fusion and reduced synaptic facilitation. This suggests that TDP-43 filament accumulation compromises the fusogenicity of the readily releasable pool of synaptic vesicles docked at the presynaptic active zone and/or the associated synaptic vesicle release machinery. The proximity of TDP-43 filaments to synaptic vesicle and the presynaptic active zone components revealed by proximity labelling, as well as contacts between filaments and synaptic vesicles and the plasma membrane revealed by cryo-ET, suggests that these interactions may underlie presynaptic dysfunction. Future studies could exploit the models and methods we present here to further interrogate the molecular basis of synaptic TDP-43 filament accumulation and presynaptic dysfunction, including the roles of synaptic TDP-43 filament interactions. At later timepoints, an increased frequency of spontaneous synchronous calcium transients was observed in neurons treated with TDP-43 filaments, suggesting that they resulted in network hyperexcitability. Future studies will be required to investigate if hyperexcitability is caused by altered intrinsic neuronal properties and/or altered synaptic activity, and if this results from chronic presynaptic dysfunction.

Presynaptic dysfunction and neuronal hyperexcitability are among the earliest hallmarks of TDP-43 proteinopathies^33^, including ALS and FTD, with evidence for contributions from both intrinsic neuronal hyperexcitability and altered synaptic transmission^76^. Our results suggest that presynaptic TDP-43 filament accumulation may underlie presynaptic dysfunction and neuronal hyperexcitability in disease. Synaptic TDP-43 filament accumulation may also underlie previous reports of synaptic dysfunction and neuronal hyperexcitability in mice overexpressing TDP-43 lacking a functional NLS^50,77^, as well as in patient iPSC-derived motor neurons harbouring pathogenic *TARDBP* variants^78^, which both develop cytoplasmic TDP-43 inclusions.

Knockdown of TDP-43 in human iPSC-derived cortical neurons, to mimic nuclear TDP-43 depletion resulting from pathological TDP-43 assembly, has also been shown to lead to synaptic and network dysfunction^79–81^. This appears to be largely driven by the cryptic splicing of synaptic proteins, including the presynaptic active zone complex protein unc13A. These splicing events are not conserved in mice^81–83^, so cannot account for the synaptic and network dysfunction we observed using mouse primary neurons, or that was previously observed in mice exhibiting TDP-43 inclusions^50,77^. In addition, we did not observe nuclear TDP-43 depletion in our models. This suggests that loss of nuclear TDP-43 function and TDP-43 filament gain of function might jointly interfere with synaptic functions.

TDP-43 filaments from different types of FTLD-TDP have been shown to have distinct propensities to propagate and cause toxicity in cell and mouse models^30,31,49^. We previously showed that TDP-43 filaments from different types of FTLD-TDP have distinct core folds^5,9,10^, suggesting that they might act like prion strains. Future studies could use the models and methods outlined here to investigate if distinct molecular environments and interactions of disease-characteristic TDP-43 filament folds underlie differences in their rates of propagation and toxicity.

In addition to TDP-43, pre-inclusions of tau, α-synuclein, superoxide dismutase 1 and prions have been observed at synapses in neurodegenerative diseases characterised by their assembly^84–87^. Moreover, synaptic dysfunction and network hyperexcitability are common features of these diseases^33^. This suggests that there may be shared synaptic mechanisms involving toxic gain of function of intracellular amyloid filaments across neurodegenerative diseases. The proximity labelling and cryo-ET approaches we outline here could be adapted to investigate if there are common synaptic mechanisms of intracellular filaments beyond TDP-43.

## CONCLUSION

We have shown that TDP-43 filaments accumulate at synapses upon their internalisation by cultured neurons. This was followed by synaptic and network dysfunction. Our work provides a possible link between observations of synaptic TDP-43 pre-inclusions and early synaptic dysfunction in disease. Targeting synaptic TDP-43 filaments and their interactions may represent strategies to limit TDP-43 pathology and synaptic dysfunction in neurodegenerative diseases.

## METHODS

### Human tissue samples

Frozen postmortem frontotemporal cortex tissue samples from individuals with FTLD-TDP Type A were from the Dementia Laboratory Brain Library at Indiana University School of Medicine. Neuropathological diagnosis of FTLD-TDP Type A was made according to the criteria set out in^12^. The individuals carried *GRN* variants associated with FTLD-TDP Type A and had clinical presentations of behavioural variant FTD (bvFTD). Control frozen postmortem frontotemporal cortex tissue samples from aged individuals without amyloid pathology and neurological disorders were from the Queen Square Brain Bank for Neurological Disorders at University College London Queen Square Institute of Neurology. The use of human tissue samples in this study was approved by the ethical review processes at each institution. Informed consent was obtained from the patients’ next of kin. Additional clinicopathological details are given in Supplementary Table 1.

### Extraction of pathological TDP-43 filaments

Extraction of pathological TDP-43 filaments from human post-mortem frontotemporal cortex tissue samples was performed based on^5^, with Benzonase modifications from^88^. Grey matter (125 mg) was dissected and homogenised using a Polytron (Kinematica) in 625 µL of extraction buffer (EB) consisting of 10 mM Tris-HCl, 800 mM NaCl, 1 mM ethylenediaminetetraacetic acid (EDTA), 1 mM dithiothreitol (DTT), 10% sucrose and 1X protease and phosphatase inhibitor cocktail (ThermoFisher, A32965). The homogenate was incubated for 30 min on ice, followed by sonication at 50% amplitude (Qsonica Q700) in a water-bath for 10 min, with 10 s on and 20 s off cycles. For every 625 µL of homogenate, 2.5 mL of EB without NaCl was added, along with 10 μL of Benzonase (Millipore, 3594276) and 10 μL of 100 mM MgCl_2_, to reach a final concentration of 1.25 u/μL Benzonase and 3 mM MgCl_2_. The homogenate was incubated for a further 30 min at 21°C and then brought to 1% Sarkosyl by addition of 25% Sarkosyl in water, followed by incubation at 37°C overnight with shaking at 200 RPM. The sample was then diluted 2X with EB and centrifuged at 3,000 xg for 10 min at 21°C. The supernatant was retained and centrifuged at 166,000 xg for 25 min at 25°C using a TLA55 rotor (Beckman). The resulting pellet was resuspended in 1 mL EB and centrifuged at 166,000 xg for 25 min at 25 °C. The resulting pellet was resuspended in 25 μL sterile phosphate-buffered saline (PBS) and stored at 4°C short term and -80°C long term.

For Airyscan2 confocal imaging, electrophysiology and calcium imaging, TDP-43 filaments were extracted from human post-mortem frontotemporal cortex tissue samples as described in^5^. Grey matter (100 mg) was dissected and homogenised using a Polytron (Kinematica) in 4 mL of extraction buffer (EB2) consisting of 10 mM Tris-HCl, 0.8 M NaCl, 10% sucrose, 1 mM ethylene glycol-bis(β-aminoethyl ether)-N,N,N’,N’-tetraacetic acid (EGTA) and 1X protease and phosphatase inhibitor cocktail (ThermoFisher, A32965). The homogenate was brought to 2% Sarkosyl by addition of 25% Sarkosyl in water and incubated at 37°C overnight with shaking at 200 RPM. The homogenate was then centrifuged at 27,000 xg for 10 min at 25°C. The supernatant was retained and centrifuged at 166,000 xg for 25 min at 25°C. The pellet was resuspended in EB2 containing 1% Sarkosyl and sonicated in a water bath continuously for 5 min at 50% amplitude (Qsonica Q700). Following sonication, the solution was diluted 4X in EB2 containing 1% Sarkosyl and incubated at 37°C for 1 h with shaking at 200 RPM. The solution was then centrifuged at 17,000 xg for 5 min at 25°C. The supernatant was retained and centrifuged at 166,000 xg for 20 min at 25°C. The pellet was resuspended in EB2 containing 1% Sarkosyl and incubated at 37°C for 1 h with shaking at 200 rpm, followed by centrifugation at 166,000 xg for 20 min. The resulting pellet was resuspended in phosphate-buffered saline (PBS) at 250 μL/g initial grey matter used. Control aged human tissue samples lacking amyloid pathology were extracted using the same protocols.

### Immunoblotting

For polyacrylamide gel electrophoresis, samples were incubated in 1X lithium dodecyl sulfate (LDS) sample buffer (Thermo Fisher Scientific, NP0007) containing 2.5% β-mercaptoethanol for 10 min at 95°C, then resolved using 4-12 % Bis-Tris gels (Thermo Fisher Scientific, NP0321BOX) for 45 min at 200 V. Resolved proteins were transferred onto nitrocellulose membranes using a Trans-Blot Turbo system (BioRad, 1704158) using the High Molecular Weight setting. For dot blots, native samples were diluted in 200 μl PBS and loaded directly onto nitrocellulose membranes using a 96-well dot blot apparatus connected to a vacuum pump.

Membranes were blocked for 1 h at 21°C using a blocking buffer containing 1% (w/v) BSA and 0.2% Tween-20 in PBS. The membranes were incubated with primary antibodies diluted in blocking buffer for 1 h at 21°C or overnight at 4°C. Pierce High Sensitivity Streptavidin-HRP (Thermofisher, 21130; 1:5000) was used where indicated. The membranes were washed 3 times at 10 min intervals using PBST and incubated with secondary antibodies diluted in blocking buffer for 1 h at 21°C, protected from light. The membranes were then washed 3 times at 10 min intervals using PBST. For fluorescence-conjugated secondary antibodies, the membranes were then imaged directly. For HRP-conjugated secondary antibodies, the membranes were developed using an enhanced chemiluminescence (ECL) kit (Asheram, RPN2106). Membranes were imaged using a Gel Doc (BioRad). Densitometry of do blots was performed using FIJI.

The following the primary antibodies were used: anti-phospho-serine 409 and 410 TDP-43 (Cosmo-Bio, TIPPTD-M01; 1:3000); anti-TDP-43 (Proteintech, 10782-2-AP; 1:5000); and anti-postsynaptic density protein 95 (NeuroMab, 75-028; 1:1000). The following secondary antibodies were used: Goat Anti-Rabbit IgG (H+L)-HRP Conjugate (BioRad, 176515; 1:4000); Goat Anti-Mouse IgG (H+L)-HRP Conjugate (BioRad, 176516; 1:4000); Goat Anti-rabbit IgG-Dylight 800 (Cell Signal, 5151; 1:5000); and StarBright Blue 700 Goat Anti-Mouse IgG (BioRad, 12004158; 1:4000).

### Immunogold negative-stain electron microscopy

TDP-43 filament extracts were diluted 1:3 and 3 μL was deposited on glow-discharged carbon-coated 400-mesh copper grids (Electron Microscopy Sciences, 71150). Samples were incubated for 40 s and then blotted with grade 1 Whatman filter paper (Whatman, 1001-070). The grids were then blocked using 0.1% (w/v) fish skin gelatin in PBS for 10 min at 21°C. The grids were then blotted and incubated with anti-TDP-43 primary antibody (Proteintech, 10782-2-AP; 1:25) diluted in blocking buffer for 3 h at 21°C. The grids were blotted and washed once with 1 mL of blocking buffer, followed by incubation with gold-conjugated goat anti-rabbit secondary antibody (Sigma-Aldrich, G7277, 1:20) diluted in blocking buffer for 1 h at 21°C. The grids were washed once more using 1 mL of distilled water and stained with 2 % uranyl acetate in water for 40 s, followed by blotting and air-drying for at least 20 min before imaging using an FEI Spirit transmission electron microscope at 120 keV.

### Mouse primary neurons

All experimental procedures were performed in accordance with the UK Animal (Scientific Procedures) Act 1986, under Project and Personal License authority, and was approved by the Animal Welfare and Ethical Review Bodies at the MRC Laboratory of Molecular Biology and the University of Edinburgh. All mice were housed with unlimited access to food and water on either a 12 h (CD1 mice) or 14 h (C57BL/6J mice) light-dark cycle at room temperature (20-22°C) and 45-65% humidity. Mice were killed by Schedule 1 procedures in accordance with UK Home Office Guidelines. Specifically, pregnant adult (> 2 months old) female mice were killed by exposure to rising CO_2_ concentrations, followed by decapitation to confirm death. Embryonic day (E) 16.5 to 18.5 embryos were killed by decapitation, followed by removal of the brain. Primary cortical and hippocampal cultures were prepared from these mouse embryos immediately after culling.

Primary mouse cortical and hippocampal neurons were isolated from embryonic (E) day 18.5 CD1 wild-type mice based on^89^. Cortices and hippocampi were dissected in ice-cold HBSS (Ca2+ and Mg2+ free; Gibco, cat. no. 14175095) containing 0.11 mg/mL sodium pyruvate (Gibco, cat. no. 12539059), 0.1% glucose and 10 mM HEPES (Gibco, 15630056) and dissociated for 20 min at 37°C with trypsin (0.25% w/v; Gibco, 15090-046). For plating, neurons were resuspended in plating medium containing 86.55% MEM (Gibco, 21090022), 10% heat-inactivated FBS (Gibco, 11573397), 0.45% glucose, 1 mM sodium pyruvate and 2 mM GlutaMax (Gibco, 35050038). For proximity labelling and immunoblotting, neurons were plated on poly-L-lysine-coated (0.02 mg/mL) 6-well plates, at a density of 750,000 per well. For immunofluorescence imaging and electrophysiology, neurons were plated on poly-L-lysine-coated (0.02 mg/mL) 13 mm-diameter glass coverslips in 24-well plates, at a density of 120,000 per well. For calcium imaging, neurons were plated on 96-well imaging plates pre-coated with poly-L-lysine (Greiner, 655936), at a density of 24,000 per well. 4 hours after plating, neurons were maintained in Neurobasal Plus medium (ThermoFisher, A3582901) containing 1:50 (v/v) B-27 Plus supplement (ThermoFisher 175040-44) at 37 °C with 5% CO_2_. Half of the culture medium was exchanged once per week with pre-warmed fresh medium. Three days after plating, cytosine arabinoside (Ara-C) was added to the culture medium to a final concentration of 2.8 μM reduce the growth of non-neuronal cells. Neurons were matured for 14 d before use.

For SypHy experiments, primary mouse cortical neurons were isolated from E16.5-E17.5 C57BL/6J wild-type mice. Cortices were dissected, dissociated in papain for 20 minutes (10.5 U/ml; Worthington Biochemical; #LK003178) and triturated using fire-polished glass Pasteur pipettes in DMEM /F-12 (Invitrogen; #21331-020) supplemented with 10% (v/v) FBS (BioSera; #S1810-500). After a low-speed centrifugation, the tissue was resuspended in Neurobasal medium (Invitrogen; #21103-049) supplemented with 0.5 mM L-glutamine (Invitrogen; #25030-024) and 1% (v/v) B-27 supplement (50×, serum-free; Invitrogen; #17504044). Cortical neurons were plated on 25 mm glass coverslips (VWR®, Cover Glasses, Round, 631-1584), that had been previously coated with poly-D-lysine (PLL, Sigma-Aldrich, #P2636-100MG) overnight at 4°C. Coverslip-attached cells were maintained in supplemented Neurobasal medium in a humidified incubator 37°C/5% CO_2_, with cytosine β-D-arabinofuranoside (Sigma-Aldrich; #C1768) added on day *in vitro* (DIV) 3 at a 1 µM concentration to prevent glial growth.

### Human embryonic stem cell-derived cortical neurons

We used the H9 female human embryonic stem cell (hESC) line (WiCELL), previously engineered to harbour a doxycycline-inducible neurogenin-2 (NGN2) transcription factor cassette in the AAVS1 locus^55^. The hESCs were thawed and plated on Matrigel-coated (Corning, 35620) 6-well plates in modified TeSR (mTeSR) Plus medium (Stemcell Technologies, 05825) containing 1X CloneR (Stemcell Technologies 05888) and 100 U/mL Penicilin-Streptomycin (PenStrep) (ThermoFisher 15140-122). Three passages were performed before induction. For induction, hESC colonies were dissociated by incubation with pre-warmed Accutase (Gibco, A1110501) for 6 min at 37°C, then pipetted to generate a single cell suspension. The dissociated hESCs were plated on Matrigel-coated 6-well plates at a density of 500,000/well in mTeSR Plus medium containing 1% PenStrep. After 24 h, the culture medium was exchanged for induction medium consisting of Dulbecco’s Modified Eagle Medium (DMEM)/F12 (Gibco, 113300-32) with 1X Glutamax (Gibco, 35050061), 1X Minimal Essential Medium (MEM) containing non-essential amino acids (Gibco, 11140-50), 1X N2 supplement (ThermoFisher 175020-48), 1% PenStrip and 1 μg/mL doxycycline (Sigma, D9891). After 2 d, the cells were split to obtain accurate cell density measurements. Wells were washed with DMEM/F12 once prior to use. The cells were washed once with 1 ml PBS and dissociated with 1 mL Accutase for 3 min. The cells were then suspended by flowing 1 ml of DMEM/F12 over the cells, which was repeated 3 to 4 times. The resuspended cells were transferred to 15 mL Falcon tubes with a 1 mL cushion of 4% BSA. The tube was then centrifuged at 1,500 RPM for 3 min. The pelleted cells were resuspended in 300 μL neuronal maintenance media consisting of Neurobasal medium (ThermoFisher 211030-49) with 1% PenStrip, 1X Glutamax, 1X B-27 Plus (ThermoFisher 175040-44), 10 ng/mL human recombinant brain-derived neurotrophic factor (BDNF) (Peprotech, 450-02), 10 ng/mL recombinant human neurotrophin-3 (NT3) (Peprotech, 450-03) and 1 μg/mL doxycycline. Cells were plated on poly-L-Lysine (50 μg/mL) and Matrigel coated 13mm-diameter coverslips in 24-well plates or 6-well plates at densities of 100,000 and 750,000 per well, respectively. After 7 d, induced neurons were maintained in neuronal maintenance media without doxycycline. Neurons were cultured at 37 °C with 5% CO_2_. One quarter of the culture medium was exchanged twice per week with pre-warmed fresh medium. Neurons were matured for 11-14 d before use.

### Incubation of neurons with TDP-43 filaments

For immunostaining, immunoblotting, antibody-targeted proximity labelling and calcium imaging, TDP-43 filament extracts (or control extracts) were used at a dilution of 2 μL/mL culture medium. For Airyscan2 confocal imaging, this was reduced to 1 μL/mL culture medium to reduce background signal. For SypHy imaging and electrophysiology, TDP-43 filament extracts (or control extracts) were used at a dilution of 4 μL/mL culture medium.

### Immunofluorescence imaging of cultured neurons

Neurons grown on 13mm-diameter coverslips were fixed using 4% (w/v) paraformaldehyde (PFA) for 20 min at 4 °C, followed by 10 min at 21°C. They were then washed 3 times with PBS at 10 min intervals and permeabilised using 0.5% Triton-X100 in PBS for 15 min at 21 °C. The permeabilised neurons were blocked using 1% (w/v) bovine serum albumin (BSA) in PBS containing 0.2% Tween-20 for 1 h at 21°C. They were then incubated overnight at 4 °C with primary antibodies diluted in either PBS or blocking buffer. Streptavidin Alexa Fluor 488 conjugate (Thermo Fischer Scientific, S11223; 1:2000) was used where indicated. The neurons were then washed 3 times with PBS at 10 min intervals and incubated for 1 h at 21°C with fluorescence-conjugated secondary antibodies diluted in either PBS or blocking buffer, protected from light. The neurons were then washed 3 times with PBS at 10 min intervals, followed by counter-staining using Hoechst at 5 μg/mL.

For validation of colocalisation between TDP-43 filaments and proximal proteins (ED Figure 5), neurons were permeabilised using 0.1% saponin and 1% BSA in PBS for 1 h at 21 °C, washed using PBS containing 0.01% saponin and incubated with antibodies in PBS containing 0.01% saponin and 0.1% BSA. For pre- and post-permeabilisation experiments, neurons were blocked using 1% BSA in PBS for 1 h at 21 °C directly following fixation. Antibody staining then proceeded as described, with antibodies diluted in PBS. The neurons were then permeabilised using 0.5% Triton-X100 in PBS for 15 min at room temperature, followed by additional blocking using 1% (w/v) BSA in PBS containing 0.2% Tween-20 for 1 h at 21°C. The neurons were than incubated with primary antibody overnight at 4 °C, followed by 3 washes using PBS and incubation with secondary antibodies for 1 h at room temperature. The neurons were washed 3 times using PBS and counterstained using Hoechst.

Coverslips were mounted on glass slides using Prolong Glass Antifade (Thermo Fisher Scientific, P36982). The neurons were imaged using a Zeiss LSM 710 or 900 confocal microscope with 1.3 NA oil-immersion 40X objective or 1.4 NA oil-immersion 60X objective. To quantify the amount of pTDP-43 filaments associating with neurons, neuronal cytoskeleton channel was thresholded using in-built function Li from FIJI^90^, to create a mask. The pTDP-43 filaments channel was thresholded using in-built function Ostsu. The area of the pTDP-43 filaments signal within the mask was analysed using Analyze particles function in FIJI. To quantify the colocalisation of pTDP-43 with bassoon and PSD95, AiryscanZeiss 2 post-processing was performed, followed by laser shift correction. Laser shifts were calculated using 0.1 μm diameter TetraSpeck fluorescent microspheres (Thermo Fisher Scientific, T7279). We then used a de-novo macro that first masked the bassoon and PSD95 signal in 3D space and then measured the proximity of pTDP-43 signal.

The following primary antibodies were used: Anti-phospho-serine 409 and 410 TDP-43 (Cosmo-Bio, TIP-PTD-M01; 1:1,000); anti-mouse TDP-43 residues S379-S387 (in-house, 1:400); anti-class III β-tubulin (Biolegend, 801202; 1:200); anti-class III β-tubulin (Abcam, ab6046, 1:200); anti-bassoon (Merk, ABN255; 1:250); anti-postsynaptic density protein 95 (NeuroMab, 75028; 1:1,000); anti-microtubule-associated protein 2 (Abcam, ab5392; 1:500); anti-neuronal nuclei (Abcam, EPR12763; 1:200); anti-early endosome antigen 1 (BD Biosciences, 610457; 1:200); anti-clathrin (Abcam, ab21647; 1:200), anti-hook microtubule tethering protein 3 (Sigma-Aldrich, SAB1401891; 1:100); anti-chaperonin-containing T-complex subunit 5 (Proteintech, 11603-1-AP; 1:100); anti-DnaJ homolog subfamily A member 2 (Proteintech, 12236-1-AP; 1:200); anti-14-3-3 (Origene, TA327381; 1:100); anti-microtubule associated protein 1 light chain 3B (Cosmo-bio, CACCTB-LC3-2-IC; 1:50); and anti-proteasome 20S subunit β4 PSMB4 (ThermoFisher, A303-819A; 1:50). The following secondary antibodies were used, all at a dilution of 1:1,000: Goat anti-mouse IgG1 Alexa Fluor 594 (Thermo Fisher Scientific, A-21125); goat anti-mouse IgG1 Alexa Fluor 647 (Thermo Fisher Scientific, A-21240); goat anti-rabbit IgG (H+L) Alexa Fluor 488 (Thermo Fischer Scientific, A-11008); goat anti-rabbit IgG (H+L) Alexa Fluor 647 (Thermo Fischer Scientific, A-21244); goat anti-mouse IgG2a Alexa Fluor 488 (Thermo Fisher Scientific, A-21131); and goat anti-chicken IgY (H+L) Alexa Fluor 405 (Thermo Fisher Scientific, A48260;).

### Antibody-targeted proximity labelling

The protocol was adapted from^91^. Neurons grown in 6-well plates were fixed using 4% (w/v) PFA in PBS for 20 min at 4 °C, followed by 10 min at 21°C. Fixed neurons were washed once with PBST and endogenous peroxidase activity was quenched using 0.5% H_2_O_2_ for 10 min at 21°C, followed by 2 additional washes using PBST. The neurons were then permeabilised for 7 min using PBS containing 0.5% Triton-X, followed by a quick wash using PBST at 21°C. The neurons were blocked using 1% BSA in PBST for 2 h at 21°C on a rocker. Neurons were then incubated with a primary antibody against pS409/410 TDP-43 (1:1000, Cosmo-Bio TIP-PTD-M01) in blocking buffer overnight at 4°C. Following three washes with PBST at 20 min intervals, the neurons were incubated with poly-HRP-conjugated goat anti-mouse secondary antibody (ThermoFisher, B40911) in PBST containing 1% BSA for 3 h at 21°C. Neurons were then washed six times with PBST at 20 min intervals. The neurons were then incubated with 500 μL reaction buffer containing H_2_O_2_ and biotin-phenol from the Biotin XX Tyramide SuperBoost Kit (ThermoFisher, B40911) for 5 min at 21°C, to allow the HRP to catalysed the conversion of the biotin-phenol to biotin-phenoxyl radicals in the presence of H_2_O_2_, thereby biotinylating proximal biomolecules. This reaction was quenched after 5 min by the addition of 3 ml of 500 mM sodium ascorbate per well.

### Affinity purification of biotinylated proteins

Following proximity labelled, 3 million neurons were lifted from the wells using a cell scraper and lysed in 100 µL PBST containing 1.5% sodium dodecyl sulfate (SDS) and 1% sodium deoxycholate. The lysates were heated to 99°C for 1 h with mild shaking to reverse PFA cross-linking and denature proteins. The lysates were then centrifuged at 21,000 xg for 5 min at 21°C. The supernatants were brought to 1 mL with PBST and incubated with 50 μL pre-washed streptavidin beads from the Pierce MS-Compatible Magnetic IP Kit (ThermoFisher, 90408), following the manufacturer’s protocol. Briefly, the beads were sequentially washed three times with buffer A, two times with buffer B, once with PBST, twice with PBST with 1 M NaCl and twice with PBS. The beads were collected and resuspended in 100 μL PBS.

### Mass spectrometry sample preparation

Proteins bound to beads were resuspended in 20 mM HEPES containing 2 mM dithiothreitol (DTT), 2 M urea and 5 ng/μL sequencing grade trypsin (Promega) an incubated for 3 h at 25°C. The beads were collected and the flow through was transferred to clean tubes. The beads were washed sequentially, once with 2 M urea in 20 mM HEPES and once with 1 M urea in 20 mM HEPES, and the flow throughs combined. The resulting samples were alkylated with 4 mM iodoacetamide (IAA) in the dark for 30 min at 25°C. An additional 0.1 μg of trypsin (Promega) was added to the samples, which were digested overnight at 25°C. The next day, samples were acidified with formic acid (FA) and desalted using C18 stage tips (3M Empore) packed with poros oligo R3 resin (Thermo Scientific). The stage tips were equilibrated with 80% acetonitrile (MeCN) in 0.5% FA, followed by 0.5% FA. Bound peptides were eluted step-wise with increasing MeCN concentrations at 30, 50 and 80%, and eluates were partially dried using a Speed Vac (Savant) before LC-MS/MS analysis.

### Mass spectrometry data acquisition

Mass spectrometry data acquisition was carried out using a Q Exactive HF-X mass spectrometer (Thermo Fisher Scientific) equipped with an Ultimate 3000 RSLC Nano system (Thermo Fisher Scientific). Peptides were separated on an EASY-Spray 50 cm x 75 μm ID, PepMap RSLC C18 2 μm column, using buffer A (5% dimethylsulfoxide (DMSO), 95% water and 0.1% FA) and buffer B (5% DMSO, 75% MeCN, 20% water and 0.1% FA) and eluted at 250 nL/min flow rate using an increasing MeCN gradient. The mass spectrometer, which was operated in data-dependent acquisition (DDA) mode, performed full-scan MS1 acquisitions at m/z = 380-1750, with a resolution of 120 K, followed by MS2 acquisitions of the 30 most intense ions with a resolution of 15K and normalized collision energy (NCE) of 27%. Dynamic exclusion was set at 50 s.

### Mass spectrometry data analysis

The raw data from LC-MS/MS were processed using MaxQuant (Cox and Mann, 2008) using the integrated Andromeda search engine (v.1.6.17.0 or v 2.4.2.0). Data were searched against the UniProt_H sapiens_reviewed (downloaded in December 2020) or UniProt_M musculus_reviewed (downloaded in November 2020) protein databases. Carbamidomethylation of cysteines was set as fixed modification, while methionine oxidation and protein N-terminal acetylation were set as variable modifications. Tryptic peptides with a maximum of two missed cleavage sites were searched. Protein quantification requirements were set at 1 unique and 1 razor peptide. The search feature “match between runs” was not selected. Other parameters in MaxQuant were kept as default values. Protein abundances were normalized using the MaxLFQ label-free normalization algorithm built into MaxQuant. The MaxQuant output file, proteinGroups.txt, was then analyzed using the Perseus software (v 1.6.15.0 or v 2.0.7.0). After uploading the matrix, the data was filtered to remove proteins identified from a reverse database (decoy). Proteins identified with modified peptides only and common contaminants such as organic solvent clusters were filtered out. Datasets were exported as .txt files for further analysis.

The subsequent analysis pipeline was built using the Python package Pandas. The tables for the 1 d, 3 d and - neuron datasets were outer merged based on gene names, which combined all the rows for left and right data frames with NaN when there were no matched values in rows. Proteins that did not have gene names were removed from the analysis. Only proteins that were identified at least twice in any condition were retained for further analysis. The intensity of signal was log2 transformed and histograms were plotted to examine the distribution of the data (Supplementary Information 1). A classifier was built to filter proteins that were only identified in the experimental groups but not in the negative control groups. These proteins were used as references for future quality checks. Imputation was performed by first tabulating the mean and standard deviation of the group. Then a normal distribution was generated with a mean equal to the group mean minus 1.8 and a standard deviation equal to the group standard deviation multiplied by 0.3. The average of five randomly selected values from this distribution was used to further minimize deviation. Histograms were plotted for the imputated dataframes to ensure the imputated values were in the lower region of the plots. Principal component analysis (PCA) was performed using the Python Scikit-learn package^92^ (Supplementary Information 2). The number of variables was set to the total number of proteins and the dimensionality was reduced to 2. The first and second principle components were calculated for each condition and plotted as scatter plots. The total variance explained in the first and second principle components was also recorded.

Three subsequent data analysis pipelines were followed, which are summarised in (Supplementary Information 3). In the first analysis, 2-tailed Student’s t-tests were performed between the experimental group and the negative control groups. Proteins were selected if they exhibited positive fold-changes and p-values <0.05 when the positive group was compared to both negative control groups. The same analysis was carried out for proteins identified in the absence of neurons. The lists of selected proteins in the presence and absence of neurons were compared. Proteins that were selected in the presence of neurons, but not observed in the absence of neurons, were retained (Supplementary Data Figures 4 and 5). In the second analysis, 2-tailed Student’s t-tests were performed to compare the groups in the presence and absence of neurons. Proteins were selected if they exhibited a positive fold-change in the presence of neurons compared to the absence of neurons and p-values < 0.001 for proteins identified in the presence of neurons compared to the absence of neurons for any of the groups (experimental and controls). 2-tailed Student’s t-tests were then performed to compare these filtered proteins in the experimental groups to the negative control groups. The proteins that had positive fold-changes and p-values <0.05 were retained (Supplementary Information 6). In the third analysis, 2-tailed Student’s t-tests were performed to compare the experimental group in the presence of neurons to all groups (experimental and controls) in the absence of neurons. Proteins were selected if they exhibited positive fold changes and p-values < 0.001 when the experimental group in the presence of neurons was compared to all groups in the absence of neurons. 2-tailed Student’s t-tests were then performed to compare these filtered proteins in the experimental group with both negative control groups in the presence of neurons. The proteins that had positive fold changes and p-values <0.05 were retained (Supplementary Information 7 and 8). The common proteins between the first and third analyses were selected as the final list of significantly enriched proximal proteins (Supplementary Data). The codes used for these analysis pipelines were uploaded in GitHub under the link: https://gitfront.io/r/HeidyChen/DjoUQ5rPDeTW/ProximityLabelling/.

### Gene Ontology enrichment analysis

Gene Ontology (GO) enrichment analysis was performed using the PANTHER Classification System (Mi et al., 2013). The significantly enriched proximal proteins from the 1 d and 3 d mouse and human datasets were analysed separately. Statistical over-representation tests for Biological Process and Cellular Component terms were performed. The reference lists for the mouse and human datasets were set to the reviewed mouse and human brain genome from UniProt, respectively (both downloaded in September 2022). Fisher’s Exact test was used with the Bonferroni correction for multiple comparisons. Only terms with p-values < 0.05 after correction for multiple comparisons were retained (Supplementary Data).

### Synaptosome isolation from primary mouse cortical neurons

The synaptosome isolation protocol was adapted from^93^. Neurons were lifted from 4 wells of a 6-well plate in 100 μL dissociation buffer consisting of 10 mM HEPES pH7.4, 2 mM EDTA, and 1X protease and phosphatase inhibitor cocktail (ThermoFisher, A32965) using a cell scraper. The lifted neurons were transferred to a Douncer and dissociated with 7 strokes. The dissociated neurons were centrifuged at 500 xg for 5 min at 4°C. The pellet was retained as the nuclear fraction. The supernatant was centrifuged at 10,000 xg for 15 min at 4°C. The supernatant was retained as the cytosolic fraction. The pellet, containing crude synaptosomes, was resuspended in 15 μL saline buffer containing 10 mM HEPES pH7.4, 140 mM NaCl, 5 mM KCl, 5 mM NaHCO3, 1.2 mM NaH2PO4, 1 mM MgCl2 and 10 mM glucose.

### Cryo-ET

Resuspended synaptosomes from primary mouse cortical neurons were applied to glow-discharged 2/2 µm holey carbon-coated 200 mesh gold grids (Quantifoil) and plunge-frozen in liquid ethane using a Vitrobot Mark IV (Thermo Fisher). Tilt series were acquired using a Titan Krios G4 (Thermo Fisher) electron microscope equipped with a cold field emission gun operated at 300 keV, a Selectris X energy-filter using a slit width of 10 eV and a Falcon 4i direct electron detector. Tilt series were acquired using Tomo5 (Thermo Fisher) at a pixel size of 2.39 Å and a target defocus range of -3 µm to -5 µm. A dose-symmetric tilt series acquisition scheme was used over a tilt range of ±40° at 2° increments, with each tilt receiving a dose of 4.09 e^−^ Å^–2^ fractionated over eight movie frames. Gain correction, motion correction, contrast transfer function estimation, tilt series alignment and tomogram reconstruction at a pixel size of 9.56 Å were performed in AreTomo3 (https://github.com/czimaginginstitute/AreTomo3). Reconstructed tomograms were denoised using the Noise2Noise^94^ machine learning algorithm in DenoisET (https://github.com/apeck12/denoiset) and visualised using IMOD^95^.

### Synaptosome isolation from human brain tissue

The synaptosome isolation protocol was adapted from^96^. 0.4 g grey matter from post-mortem frontotemporal cortex was minced in 2 mL of dissociation buffer (DB) consisting of 4 mM HEPES pH7.4, 1 mM EDTA, 0.25 mM DTT, 0.32 M sucrose and 1X Complete Ultra Mini EDTA-free protease inhibitor cocktail (Roche). The minced tissue was transferred to a Douncer and dissociated with 10-15 strokes. The dissociated tissue was transferred to a 50 mL thick-wall tube (Thermo Fisher, 3110-0500) and topped up with 8 mL of DB. The dissociated tissue was then centrifuged at 3,000 RPM in an SS-34 rotor for 10 min at 4°C. The supernatant was retained and centrifuged at 8,800 RPM for 15 min at 4°C. The pellet, containing crude synaptosomes, was re-suspended in 4 mL DB and loaded onto a discontinuous Percoll gradient consisting of 3%, 10%, 15% and 23% (v/v) Percoll in DB in 14 mL tubes (Beckman, 344060). The tubes were centrifuged at 31,000 xg in a Ti40 swinging bucket rotor for 15 min at 4°C. The interface of the 3% and 10% Percoll steps was collected as the myelin fraction. The interface of the 10% and 15% steps was collected as the synaptosome fraction. The fraction at the bottom of the gradient was collected as the mitochondria fraction. The collected fractions were washed by diluting in 30 mL saline buffer consisting of 10 mM HEPES pH7.4, 140 mM NaCl, 5 mM KCl, 5 mM NaHCO3, 1.2 mM NaH2PO4, 1 mM MgCl2 and 10 mM glucose. The diluted samples were then centrifuged at 125,000 RPM in an SS-34 rotor for 12 min at 4°C and the pellets were re-suspended in 25 μL saline buffer.

### Immunofluorescence imaging of human tissue sections

Tissue sections were deparaffinized by incubating 3 times with fresh Xylene for 5 minutes. Tissues were then incubated 3 times with fresh 100% ethanol for 3 minutes, followed by incubation with 3% H_2_O_2_ in Methanol for 15 minutes. Tissue was washed once using deionized water. Antigen retrieval was performed by heat treating the tissues in 500 ml of 10 mM sodium citrate buffer (pH 6.0) for 7 minutes under high pressure (15 psi). Tissue sections were then blocked using a blocking buffer containing 1% (w/v) BSA and 0.2% Tween-20 in PBS for 1 h at room temperature. The blocked tissue sections were incubated with primary antibodies in blocking buffer for 1 h at room temperature, followed by 3 washes in PBS. The sections were then incubated with Abberior Star Red (Abberior, STRED-1002; 1:1000) and Abberior Star Orange secondary antibodies (Abberior, STORANGE-1001; 1:1000) in blocking buffer for 1 h at room temperature and washed with distilled water. Lipofuscin autofluorescence was quenched using TrueBlack (Cell Signaling, 92401) according to manufacturer’s protocol. Briefly, the 20X TrueBlack was diluated to 1X in 70% ethanol just before usage. Each section was incubated with 200μl 1X TrueBlack for 30 seconds, followed by 3 washes in PBS. Sections were then counterstained using 2% Hoechst and mounted with rectangular glass cover-slips using Prolong Glass Antifade mountant (Thermo Fisher Scientific, P36982). Sections were imaged using and Abberior stimulated emission depletion (STED) microscope equipped with a MATRIX detector. Colocalisation was analysed using the Z-profile function in FIJI^90^.

### SypHy imaging

Cortical neurons were transfected on DIV 7 with synaptophysin-pHluorin (SypHy) using Lipofectamine 2000, at a ratio of 1 mg SypHy to 1 mL of Lipofectamine (Fisher Scientific; #11668027), as per the manufacturer’s instructions. Cortical cultures were seeded with either TDP-43 Filaments or control extracts DIV 10 before imaging analysis at 13-14 DIV. Cortical neurons were mounted on an imaging chamber bath (Warner; #RC-21BRFS), which enabled electrical stimulation (1 ms pulse width, 100 mA current output). The chamber was mounted on a Zeiss Axio Observer Z1 inverted epifluorescence microscope (Zeiss) with a EC Plan-Neofluar 40x oil immersion objective (NA 1.3) and Colibri 7 LED light source (Zeiss). Imaging buffer (119 mM NaCl, 2.5 mM KCl, 2 mM CaCl_2_, 2 mM MgCl_2_, 25 mM HEPES and 30 mM glucose, pH 7.4) was supplemented with 10 μM CNQX (Abcam; #ab120271) and 50 μM DL-2-amino-5-phosphonopentanoic acid (AP5; Abcam; #ab120044) and perfused continuously across the cell. Images were acquired at 2-second intervals using a Zeiss AxioCam 506 camera controlled by ZEISS ZEN 2 software. Neurons were illuminated using an EGFP filter set (exciter 470/40, emitter 525/50, dichroic 495 LP, Chroma Technology Corp). Neurons were stimulated with a train of 300 action potentials delivered at 10 Hz using a Digitimer D-330 system. After 3 min, the cells were perfused with a modified imaging buffer, where 50 mM NH_4_Cl was added in exchange of an equal concentration of NaCl. Experiments were performed at room temperature and from a minimum of four independent cell culture preparations.

Time-lapse experiments were analyzed using FIJI^90^. Images were aligned using the StackReg plugin with Rigid Body transformation type^97^. Fluorescence intensity was measured for the time-lapse experiment using the Time Series Analyzer plugin (https://imagej.net/ij/plugins/time-series.html). Circular regions of interest of 0.5 μm in diameter were placed on presynaptic boutons that responded to challenge with NH_4_ supplemented buffer. Only nerve terminals that responded to action potential stimulation were selected for analysis. The ΔF/F_0_ of the sypHy response was calculated for each experiment and was normalised to either the maximum fluorescence intensity during stimulation or NH_4_ buffer perfusion, respectively. A linear regression was applied to the first 6 seconds of the evoked sypHy response that was normalised to stimulation.

### Electrophysiology

Neurons were used for electrophysiological recordings 3-7 d after seeding (DIV17-21). Coverslips were perfused with an artificial CSF containing 126 mM NaCl, 3 mM KCl, 1.25 mM NaH_2_PO_4_, 26.4 mM NaHCO_3_, 10 mM glucose, 2 mM CaCl_2_, 2 mM MgSO4 and 1 µM SR-9553, saturated with 95% O_2_ and 5% CO_2_. Borosilicate glass pipettes for whole-cell recordings (1.5 mm outer diameter, 0.86 mm inner diameter, Science Products) were pulled with a P-1000 horizontal puller (Sutter Instruments) to 3-5 MΩ. Pipettes used for the presynaptic cell, to induce action potentials in current clamp, were filled with an intracellular solution containing 10 mM 4-(2-Hydroxyethyl) piperazine-1-ethane-sulfonic acid (HEPES) pH 7.25, 140 mM K-Gluconate, 2 mM MgCl_2_, 4 mM NaATP, 0.1 mM EGTA, 10 mM phosphocreatine, 0.4 mM GTP and 4 mg/ml biocytin. Pipettes for recording excitatory postsynaptic current (EPSC) in the postsynaptic cell in voltage clamp used an intracellular solution containing 10 mM HEPES pH 7.25, 125 mM CsCH_3_SO_3_, 2 mM MgCl_2_, 1 mM CaCl_2_, 4 mM NaATP, 10 mM EGTA and 0.4 mM GTP. For paired recordings, the cells were considered monosynaptically coupled if the time between the peak of the presynaptic action potential and postsynaptic evoked EPSC was <4 ms. Recordings were made using pClamp10 (Molecular Devices) with a Multiclamp 700B amplifier (Axon Instruments) and digitised using a Digidata 1440 A (Axon Instruments).

### Calcium imaging

Adeno-associated virus 2/1 (AAV2/1) particles were produced by co-transfection of AAV Pro HEK 293T cells (Takara) with pAAV2/1 (Addgene #112862), pAdDeltaF6 (Addgene #112867) and pAAV.Syn-NES-jRGECO1a.WPRE.SV40 (Addgene #100854) using polyethylenimine (1 mg/mL). Viral particles were purified by iodixanol gradient centrifugation as described^98^ and purity was confirmed by SDS-PAGE with Coomassie staining. Neurons were incubated with viral particles (3750 vg/cell) at DIV7 for 7 d. TDP-43 filaments and control brain extracts were added at DIV14 and incubated for 9-14 d. Live imaging was performed using a 20x objective on a Nikon CSU-W1 spinning disk microscope at 37°C and 5% CO_2_ at 100 ms intervals over 2 min. Out of focus images were discarded. Individual firing events were tracked using a de-novo macro in FIJI^90^.

## AUTHOR CONTRIBUTIONS

R.C. performed filament extraction, human ESC-derived neuronal culture, proximity labelling, immuno-EM, immunoblotting, immunofluorescence imaging, synaptosome fractionation, and cryo-ET data acquisition and analysis. R.C and J.P. performed mouse primary neuronal culture. J.P. performed SypHy imaging. I.S. performed electrophysiology. R.C. and S.-Y.P.-C. performed mass spectrometry and analysed mass spectrometry data. R.C. and M.H. performed calcium imaging. K.N. and B.G. identified FTLD-TDP Type A patient tissue samples and performed neuropathology. M.A.C., I.H.G. and B.R.-F. supervised the study. All authors contributed to writing the manuscript.

## ACKNOWLEDGEMENTS

We thank the individuals and their families for donating brain tissue; the Brain Library of the Dementia Laboratory at Indiana University School of Medicine for supplying tissue samples from individuals with FTLD-TDP Type A; the Queen Square Brain Bank for Neurological Disorders at University College London Queen Square Institute of Neurology, which receives support from the Reta Lila Weston Institute for Neurological Studies, for supplying tissue samples from neurologically-normal individuals without amyloid filament pathology; Mark Kotter Lab, University of Cambridge, for the provision of the ECS line harbouring a doxycycline-inducible neurogenin-2 transcription factor cassette (NGN2-OPTi-OX); A. Fellows, R. Wademan and A. Carter for help setting up ESC-derived neuronal cultures; Y. Ding, T. Stevens and S. Wingett for help with developing the mass spectrometry data analysis pipelines; F. Gao, M. McAndrew and T.S. Behr for help with cryo-ET data analysis; J. Boulanger for help with developing light microscopy data analysis scripts; T.P. Azevado for help with developing the automated calcium imaging pipeline; D. Arseni for help with IF staining of patient brain sections; staff at the MRC Laboratory of Molecular Biology Electron Microscopy, Scientific Computing, Mass Spectrometry and Light Microscopy Facilities for experimental and technical assistance; staff at the Francis Crick Institute Advanced Light Microscopy Facility for access to STED microscopy; staff at Abberior for help with STED microscopy; and D. Arseni, T.S. Behr, A. Bertolotti, A. Giblin, A. G<u>öhlke,</u> F. Hawkins, H. McMahon, S. Mishra, M. Polymenidou, S. Tetter, N. Varghese and M. Walker for discussions. This work was supported by the Medical Research Council, as part of United Kingdom Research and Innovation (also known as UK Research and Innovation) (MC_UP_1201/25 to B.R.-F. and MC_U105174197 to I.H.G.); a research collaboration between AstraZeneca UK Limited and the Medical Research Council (BSF41 to B.R.-F. and H.C.); the US National Institutes of Health (R01NS137469 to K.N. and B.R.-F; and R01-NS110437, RF1-AG071177 and R01-AG080001 to B.G.); and the Simons Foundation (529508 to M.A.C). For the purpose of open access, the MRC Laboratory of Molecular Biology has applied a CC BY public copyright licence to any Author Accepted Manuscript version arising.

