## Supplementary Figures and Tables for "Pathological TDP-43 filaments accumulate at synapses and cause synaptic dysfunction"

**Supplementary Figure 1: Estimating the concentrations of FTLD-TDP Type A patient brain-derived TDP-43 filaments. A.** Dotblot of monomeric recombinant full-length TDP-43 (Standards) and FTLD-TDP Type A patient brain-derived TDP-43 filaments at the indicated concentrations and dilutions, probed with an antibody against TDP-43. **B.** Densitometry analysis of dotblot from **(A)**, fitted to a sigmoidal curve. **C.** Concentrations of FTLD-TDP Type A patient brain-derived TDP-43 filaments extrapolated from the curve in **(B)**. Each data point represents an independent filament extraction. n=6. The mean +/- SD is shown.

**Supplementary Figure 2: Characterisation of mouse primary cortical neurons and internalisation of patient-derived pathological TDP-43 filaments by human ESC-derived cortical neurons. A.** Immunofluorescence confocal microscopy images of mouse primary cortical neurons labelled using antibodies against microtubule-associated protein 2 (MAP2; cyan), neuronal nuclei protein (NeuN; yellow) and class III β-tubulin (βIII-Tub; magenta). Scale bar, 10 µm. **B.** Representative trace of evoked excitatory postsynaptic currents from mouse primary cortical neurons. **C.** Representative trace of spontaneous excitatory postsynaptic currents from mouse primary cortical neurons. **D.** Immunofluorescence confocal microscopy images of human ESC-derived cortical neurons incubated with (+ filaments) or without (- filaments) FTLD-TDP Type A patient brain-derived TDP-43 filaments for 7 d. Neurons were labelled with an antibody against class III β-tubulin (βIII-Tub; cyan) and TDP-43 filaments were labelled using an antibody against pS409/410 TDP-43 (pTDP-43; magenta). **E.** pTDP-43 signal (white) within images masked using the βIII-tubulin signal (yellow). Signal of 0.05-3 μm$^2$ is shown in cyan. This size filter was used to exclude noise and large extracellular TDP-43 filament accumulations from quantification in **(F)**. Scale bar, 10 μm. **F.** Quantification of βIII-tubulin-masked pTDP-43 signal from **(D)** at different timepoints. The total area of pTDP-43 signal as a percentage of the βIII-tubulin signal mask is plotted. Each data point represents one technical replicate. n = 1 biological replicate. Means are shown. **G.** Immunofluorescence confocal images of mouse primary cortical neurons incubated with FTLD-TDP Type A patient brain-derived TDP-43 filaments for 1 d. TDP-43 filaments were labelled with an antibody against pTDP-43 pre- (yellow) and post- (magenta) detergent permeabilisation. Neurons were labelled with an antibody against class III β-tubulin (βIII-Tub; cyan). **H.** pTDP-43 signal (white) within images masked using the βIII-tubulin signal (yellow). Signal of 0.05-3 μm$^2$ shown in cyan. **A, D, G and H.** Scale bars, 10 μm.

**Supplementary Figure 3: Quantification of physiological nuclear TDP-43 in cultured neurons incubated with TDP-43 filaments**. **A.** Immunofluorescence confocal images of mouse primary cortical neurons (mNeuron) and human ESC-derived cortical neurons (hNeuron) incubated with (+ filaments) or without (- filaments) FTLD-TDP Type A patient brain-derived TDP-43 filaments for 7 d. Neurons were labelled with an antibody against class III β-tubulin (βIII-Tub; cyan); TDP-43 filaments were labelled using an antibody against pS409/410 TDP-43 (pTDP-43; magenta); and physiological nuclear TDP-43 in the mouse primary neurons was labelled using an antibody against mouse TDP-43 (mTDP-43; grey). Neurons were counterstained with Hoechst (yellow) to identify nuclei. Scale bar, 10 μm. **B.** Quantification of the fluorescence intensity of Hoechst-masked mTDP-43 signal from mouse primary cortical neurons shown in **(A)**, normalised to the mean mTDP-43 fluorescence intensity signal in the absence of TDP-43 filaments. Each data point represents one nucleus. Data points are colour coded by biological replicate. n = 2 biological replicates. Means +/- SD are shown. A Student's t-test was performed; ns, not significant.

**Supplementary Figure 4: Antibody-targeted proximity labelling of TDP-43 filaments in human ESC-derived cortical neurons and in the absence of neurons. A.** Immunofluorescence confocal microscopy images of human ESC-derived cortical neurons incubated with (+ filaments) or without (- filaments) FTLD-TDP Type A patient brain-derived TDP-43 filaments for 1 d, followed by antibody-targeted proximity labelling. Neurons were labelled with an antibody against class III β-tubulin (βIII-Tub; cyan), TDP-43 filaments were labelled using an antibody against pS409/410 TDP-43 (pTDP-43; magenta) and biotin was labelled using fluorescence-conjugated streptavidin (yellow). **B.** Immunoblots of affinity purified biotinylated proteins from human ESC-derived cortical neurons (beads, b) and the flowthrough (f), probed with antibodies against biotin (left) and TDP-43 (right). Neurons were incubated with (+ filaments) or without (- filaments) exogenous TDP-43 filaments, in the presence or absence (- antibody) of the antibody against pTDP-43, followed by antibody-targeted proximity labelling. **C.** Immunofluorescence confocal microscopy images of FTLD-TDP Type A patient brain-derived TDP-43 filament following antibody-targeted proximity labelling in the presence (+ antibody) or absence (- antibody) of the antibody against pTDP-43. TDP-43 filaments were labelled using an antibody against pTDP-43 (magenta) and biotin was labelled using fluorescence-conjugated streptavidin (yellow). **A and C.** Scale bars, 10 μm.

**Supplementary Figure 5: Validation of colocalisation between TDP-43 filaments and proximal proteins.** Immunofluorescence confocal microscopy images of human ESC-derived cortical neurons incubated with FTLD-TDP Type A patient brain-derived TDP-43 filaments for 3 d using antibodies against class III β-tubulin (βIII-Tub; cyan), pS409/410 TDP-43 (pTDP-43; magenta) and selected significantly-enriched proteins from antibody-targeted proximity labelling (yellow): Early endosome antigen 1 (EEA1), clathrin, hook microtubule tethering protein 3 (HOOK3), chaperonin-containing T-complex subunit 5 (TCP5), DnaJ homolog subfamily A member 2 (DNAJA2), 14-3-3, microtubule associated protein 1 light chain 3B (LC3B) and proteasome 20S subunit β4 (PSMB4). Arrows indicate examples of colocalisation. Scale bars, 10 μm.

**Supplementary Figure 6: Cryo-ET of the presynaptic cytomatrix. A-C.** Denoised tomographic slices of synaptosomes from mouse primary cortical neurons incubated with FTLD-TDP Type A patient brain-derived TDP-43 filaments for 3 d showing examples of microtubules **(A)**, F-actin **(B)** and synaptic vesicle connectors/tethers **(C)**. The yellow arrow indicates a microtubule with visible microtubule inner proteins; magenta arrows indicate TDP-43 filaments; cyan arrows indicate F-actin; and the green arrow indicates a synaptic vesicle connector/tether. Scale bars, 50 nm.

**Supplementary Figure 7: Synaptic accumulation of TDP-43 filaments in post-mortem patient brain. A.** Immunoblot of synaptosome isolation fractions (s, supernatant; cp, cell pellet; cyt, cytosolic; syn, crude synaptosomes; mye, myelin; syn+, purified synaptosomes; mit, mitochondria) from post-mortem FTLD-TDP Type A patient prefrontal cortex, probed with antibodies against postsynaptic density protein 95 (PSD95; top), pS409/410 TDP-43 (pTDP-43; middle) and TDP-43 (bottom). The arrows indicate full-length (FL) TDP-43 and C-terminal fragments (CTFs) of TDP-43. **B.** Immunofluorescence confocal microscopy image of FTLD-TDP Type A patient prefrontal cortex using antibodies against pTDP-43 (magenta) and the presynaptic active zone complex protein bassoon (yellow). Scale bar, 10 µm. **C.** Immunofluorescence stimulated emission depletion microscopy (STED) image of FTLD-TDP Type A patient prefrontal cortex using antibodies against pTDP-43 (magenta) and the presynaptic active zone complex protein bassoon (yellow). Scale bars, 1 µm and 0.5 µm, as indicated. **D.** Mean intensities of the pTDP-43 and bassoon signals from **(C)**, within the boxed region over the z sections, demonstrating colocalisation.

**Supplementary Figure 8**: **Additional analyses of SypHy imaging. A.** Relative SypHy fluorescence intensity (ΔF/F0) 160 s after termination of stimulation (equivalent to 220 s in Figure 6B). **B.** Peak relative SypHy fluorescence intensity (ΔF/F0), normalised to challenge with $NH_4$ buffer (total synaptic vesicle pool). **C.** Puncta displaying a SypHy response to action potential stimulation as a percentage of total puncta upon challenge with $NH_4$ buffer. **A, B and C.** Mouse primary cortical neurons were incubated with FTLD-TDP Type A patient brain-derived TDP-43 filaments (+ F) or with aged-matched control brain extracts (+ C) for 3 d before stimulation with a train of 300 action potentials delivered at 10 Hz and recording of SypHy fluorescence. n = 4 biological replicates. Unpaired two-tailed t-tests were performed; ns, not significant.

**Supplementary Figure 9**: **Additional analyses of paired-pulse recordings. A.** Representative post-hoc immunofluorescence confocal microscopy images of a mouse primary hippocampal neuron incubated with FTLD-TDP Type A patient brain-derived TDP-43 filaments for 3 d using antibodies against pS409/410 TDP-43 (pTDP-43; magenta) and the presynaptic active zone complex protein bassoon (yellow). The neuron was filled with biocytin during patch clamping (cyan). Scale bars, 10 µm and 1 µm, as indicated. **B.** Percentage of biocytin-masked pTDP-43 signal colocalised with bassoon, and vice versa, from **(A)** in three-dimensional space. Each data point represents one technical replicate. n= 2 biological replicates. Means +/- SD are shown. **C.** Plot of the amplitudes of the first ($1^{st}$) and second ($2^{nd}$) excitatory postsynaptic currents (EPSCs) from paired-pulse recordings of mouse primary hippocampal neurons incubated with (+F) or without (-F) FTLD-TDP Type A patient brain-derived TDP-43 filaments or with aged-matched control brain extracts (+C) for 3-4 d (d3/4) and 6-7 d (d6/7). Each data point represents one EPSC. n = 3 biological replicates. Means +/- SD are shown. **D.** Quantification of spontaneous EPSC frequency of hippocampal neurons incubated with (+F) or without (-F) FTLD-TDP Type A patient brain-derived TDP-43 filaments or with aged-matched control brain extracts (+C) for 6-7 d (d6/7). Each data point represents one recording. n = 3 biological replicates. Means +/- SD are shown. A two-way ANOVA with Tukey's multiple comparison test was performed; ns, not significant.

**Supplementary Figure 10**: **Additional analyses of calcium imaging. A.** Plot of fluorescence intensity of mouse primary cortical neurons expressing the calcium indicator jRGECO1a over 900 frames (1 frame = 100 ms). Individual spontaneous firing events were determined (green boxes) to calculate the firing frequency in Figure 6F and the average amplitude and duration of events in **(B and C)**. **B and C.** Quantification of the average amplitude **(B)** and duration **(C)** of spontaneous firing events from **(A)**. Mouse primary cortical neurons expressing the calcium indicator jRGECO1a were incubated with (+ F) or without (-F) FTLD-TDP Type A patient brain-derived TDP-43 filaments or with aged-matched control brain extracts (+ C) for 9 d (d9) and 14 d (d14). Data points are colour coded by biological replicate. n = 5 biological replicates. Means +/- SD are shown. A two-way ANOVA with Tukey's multiple comparison test was performed, *p<0.05, **p<0.01, ***p<0.001.

**Supplementary Table 1: Clinicopathological details associated with human samples used in this study.** F, female; M, male; y, years; bvFTD, behavioural-variant frontotemporal dementia; FTLD-TDP, frontotemporal lobar degeneration with TDP-43 pathology; NA, not applicable.

### Supplementary Figure 1

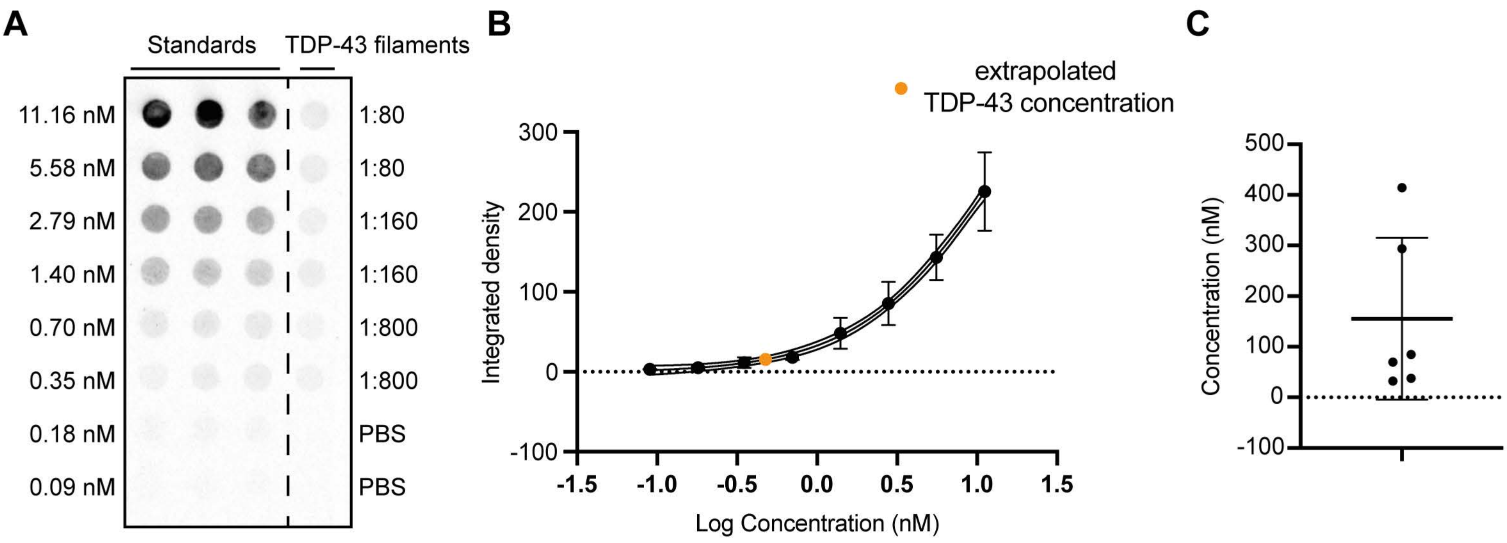

### Supplementary Figure 2

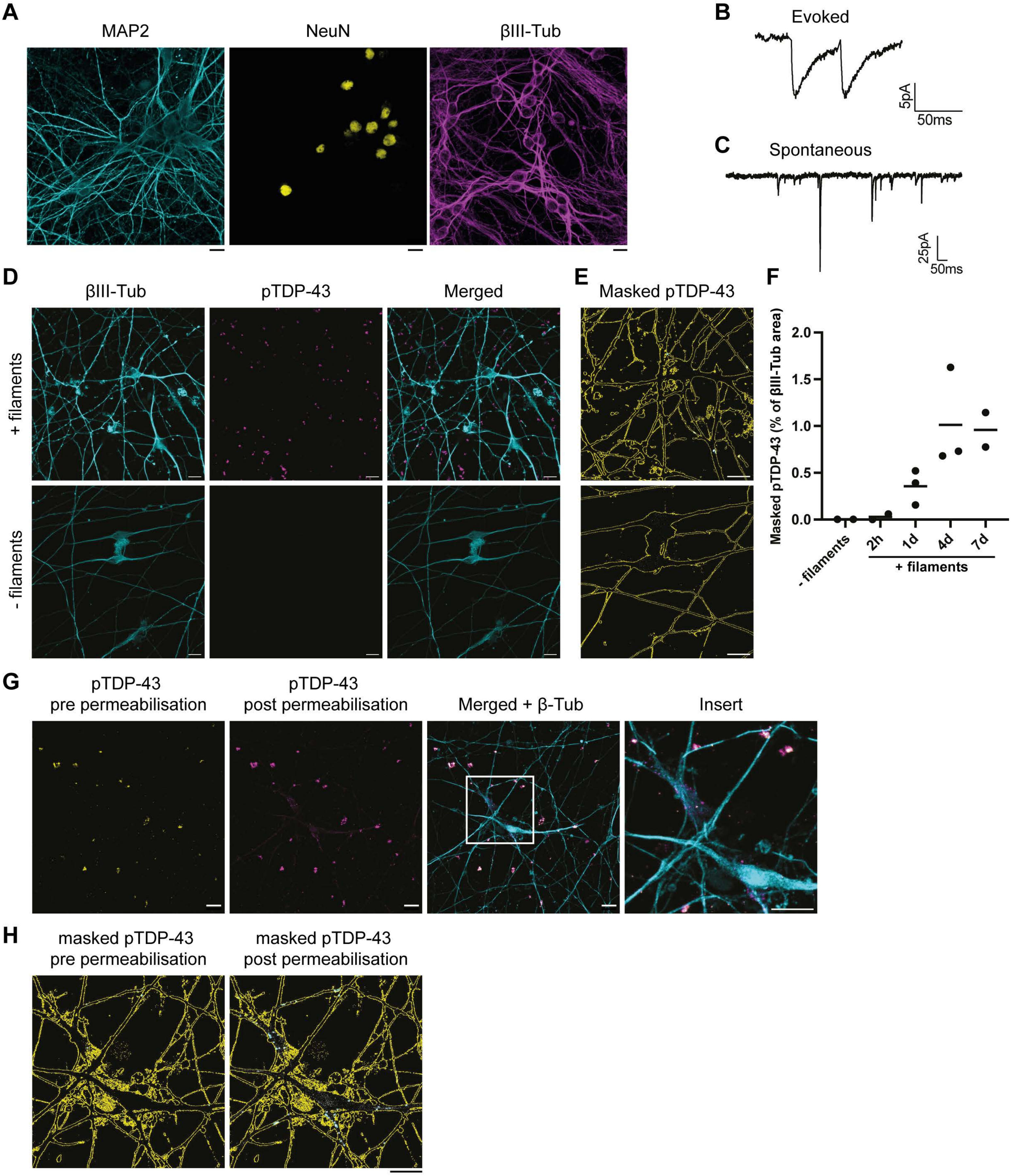

### Supplementary Figure 3

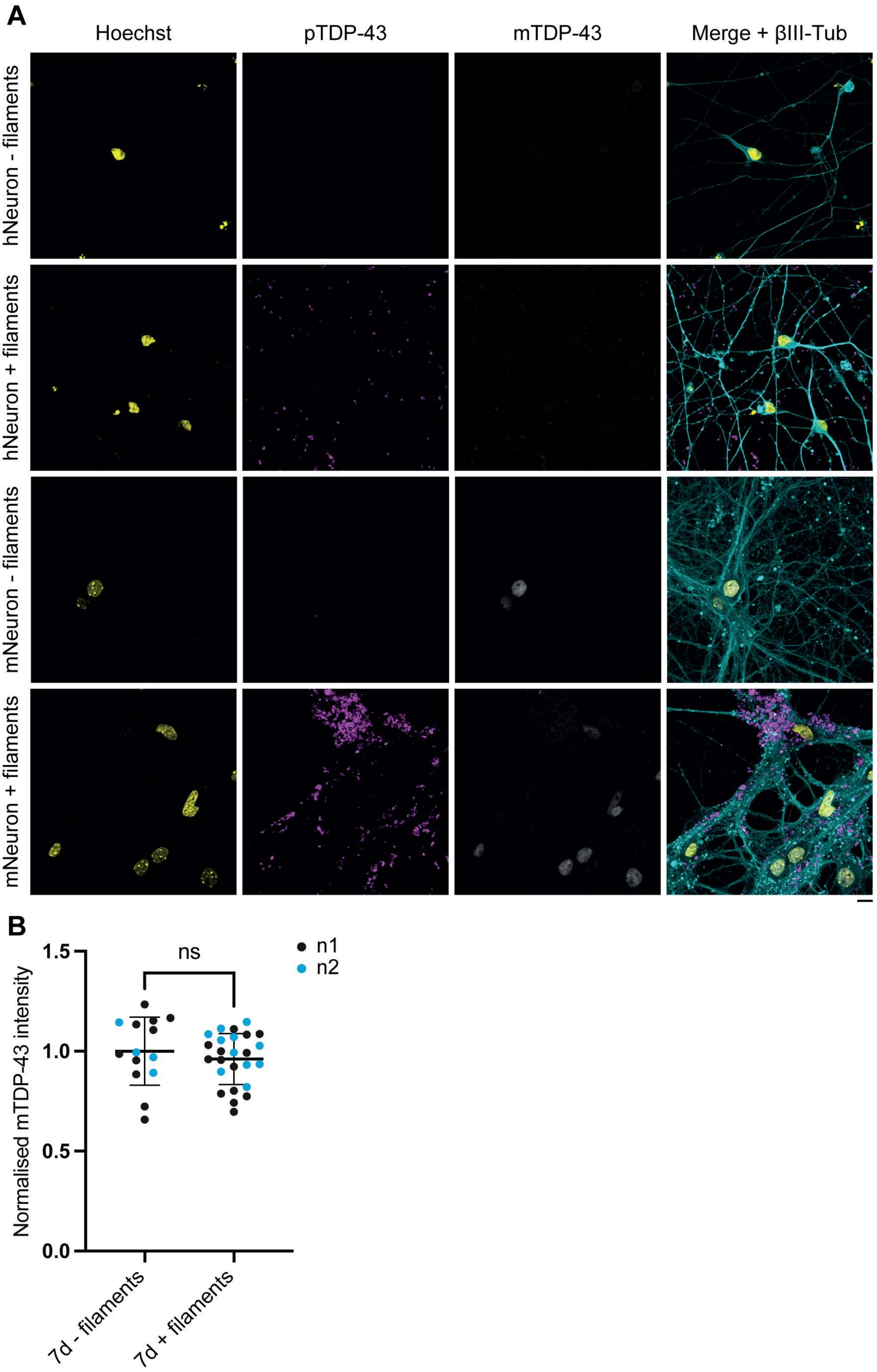

### Supplementary Figure 4

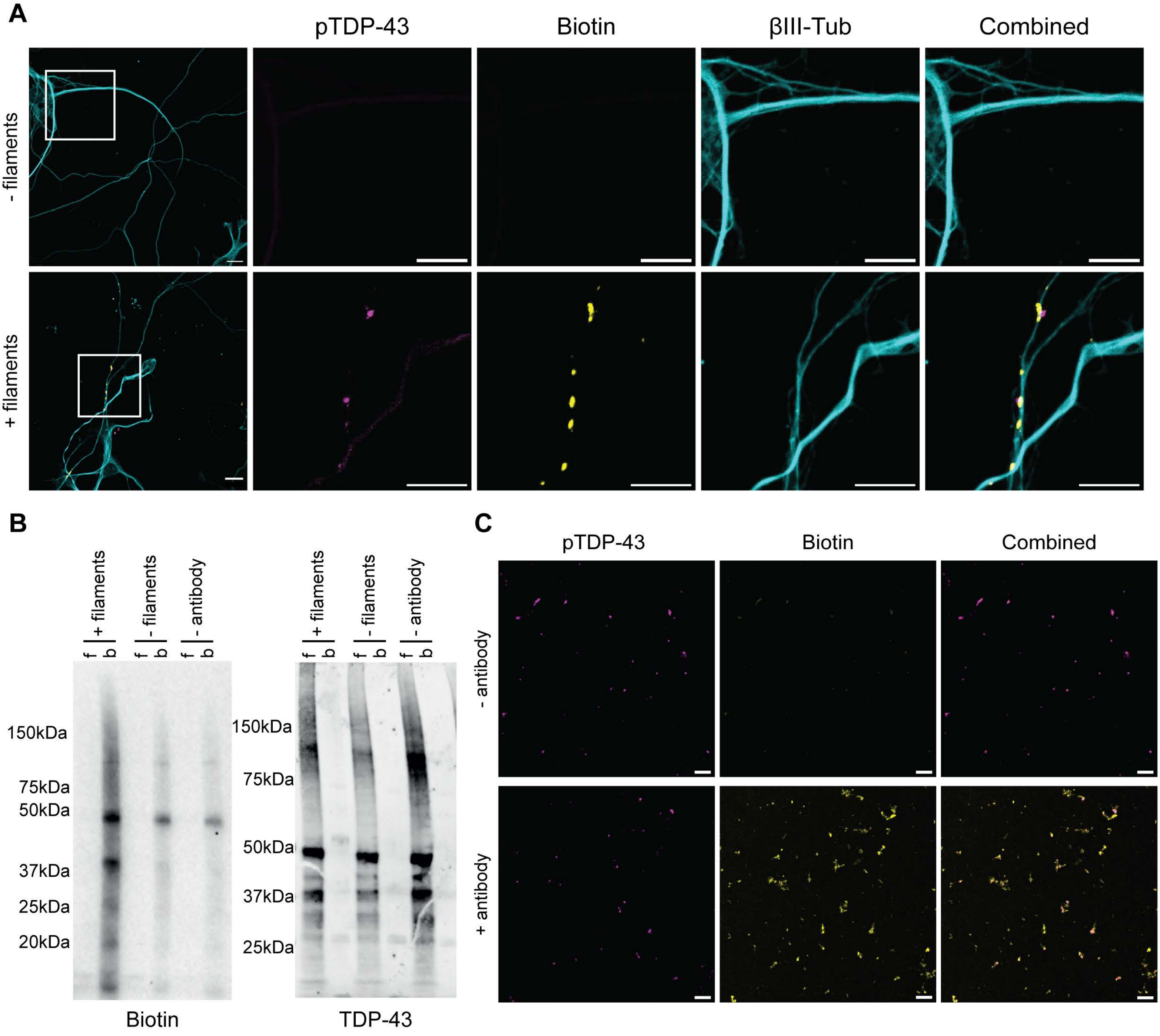

#### Supplementary Figure 5

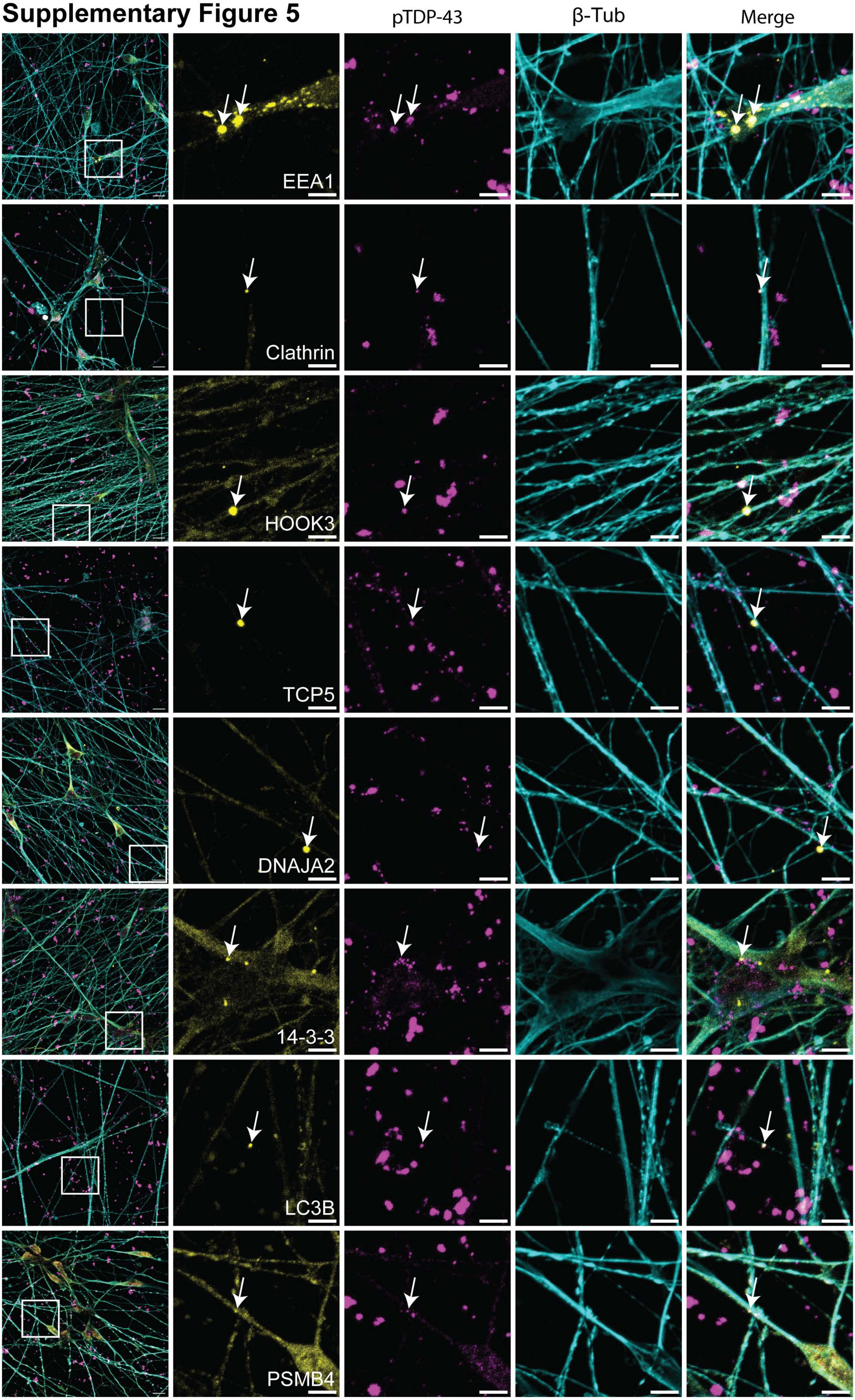

### Supplementary Figure 6

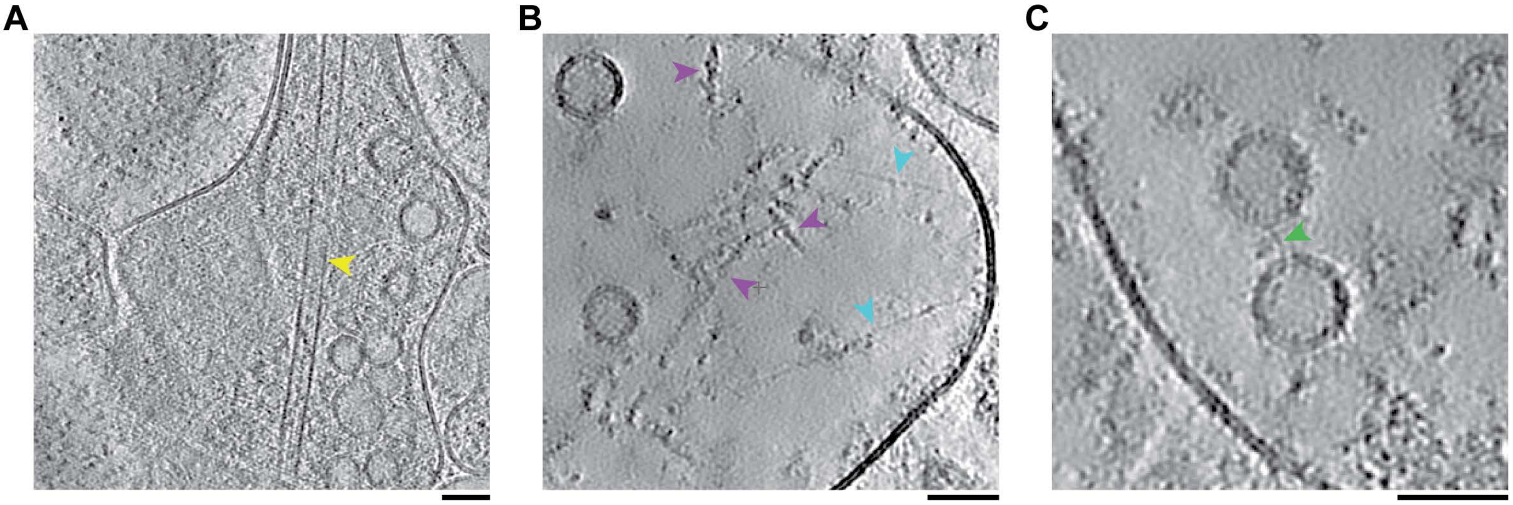

### Supplementary Figure 7

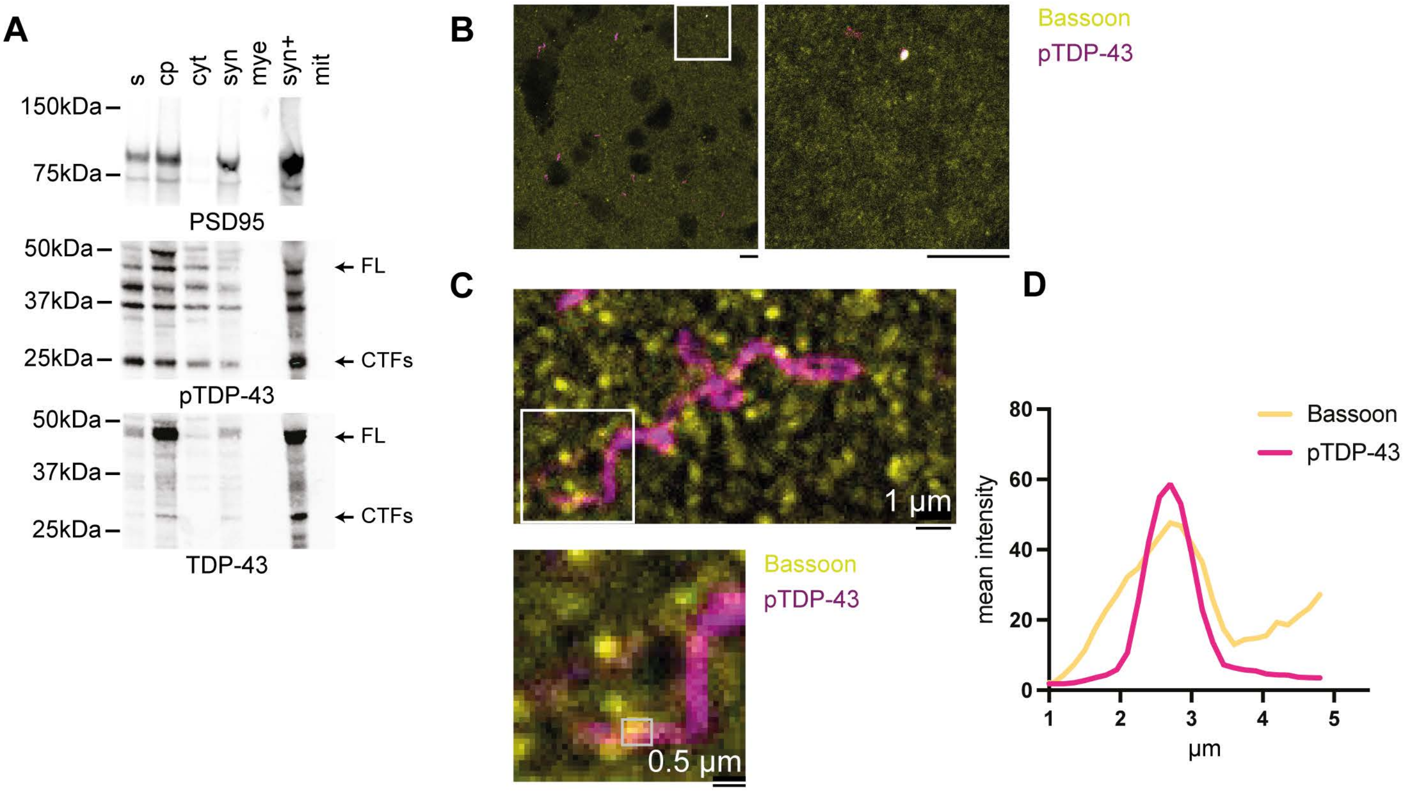

#### Supplementary Figure 8

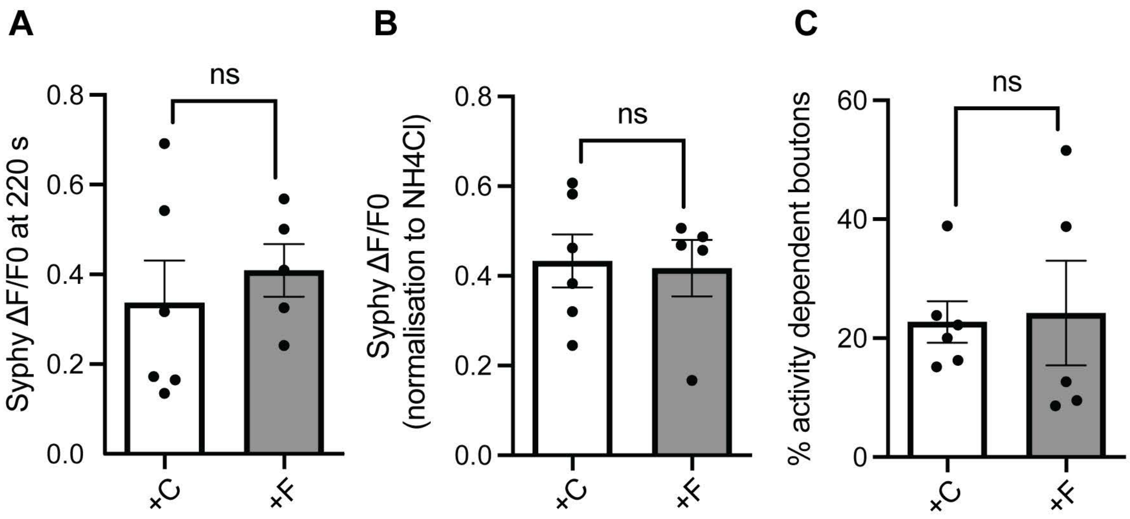

#### Supplementary Figure 9

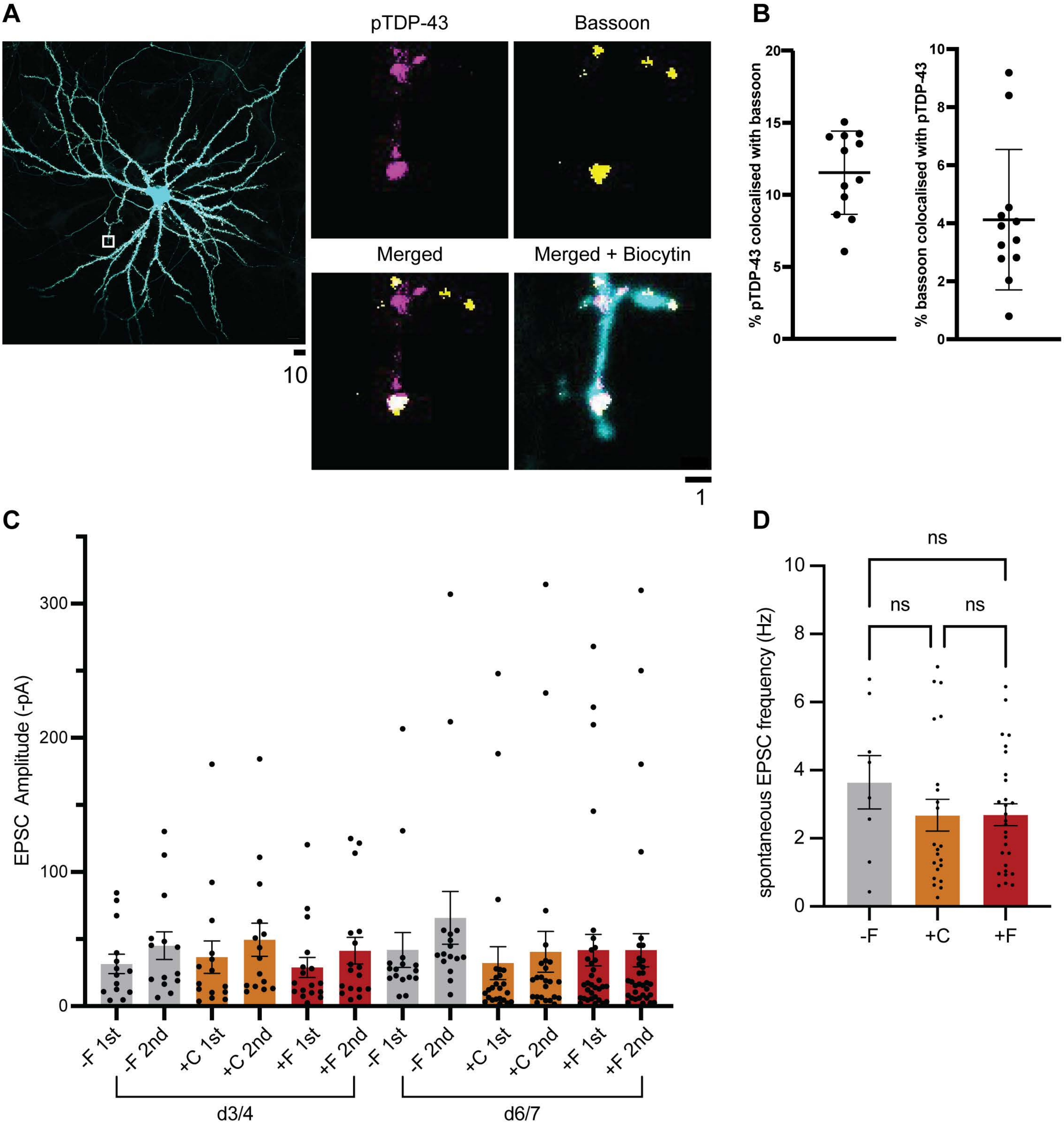

### Supplementary Figure 10

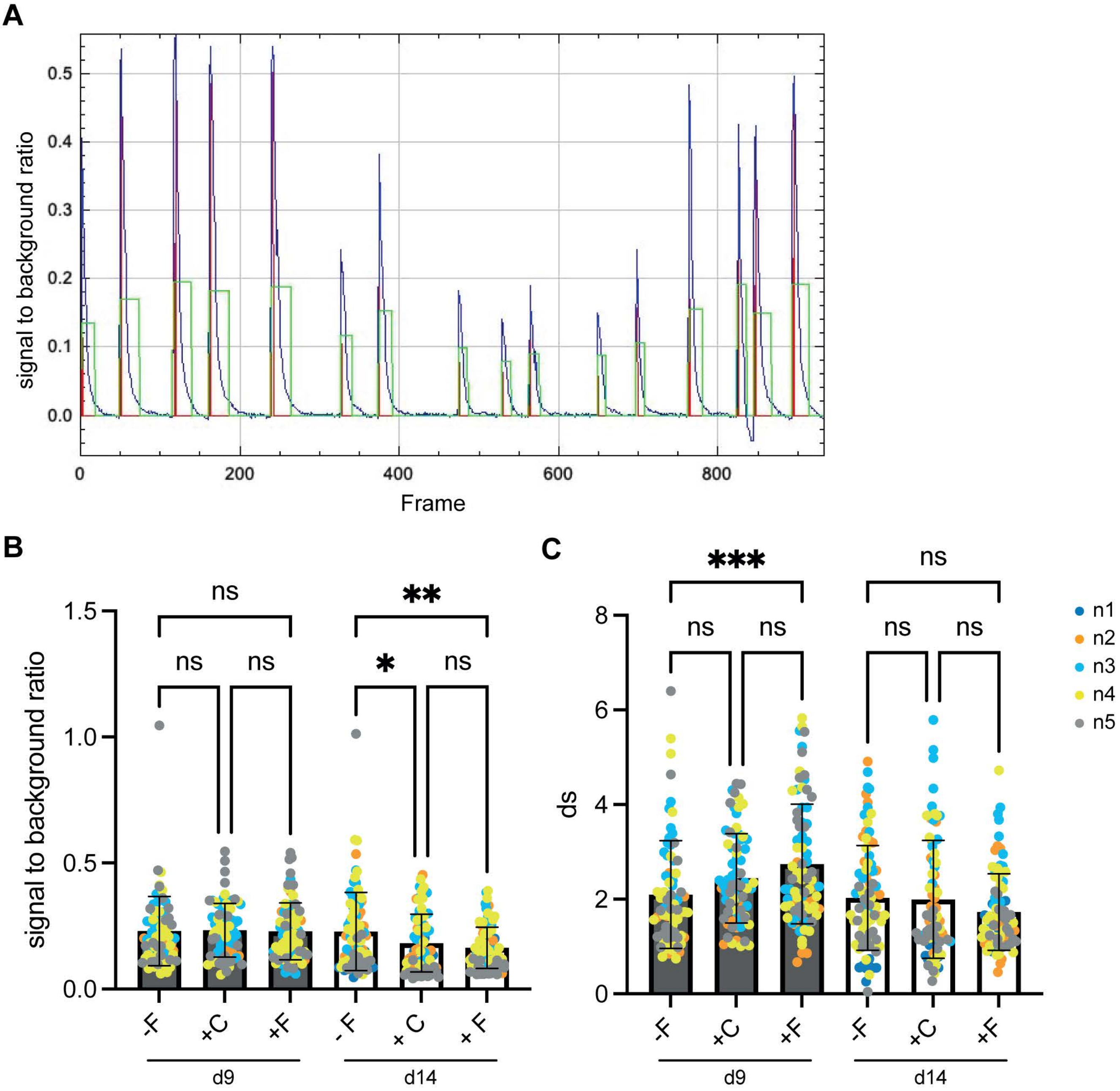

**Supplementary Table 1: Clinicopathological details associated with human samples used in this study**

| | **FTLD-TDP Type A individual 1** | **FTLD-TDP Type A individual 2** | **FTLD-TDP Type A individual 3** | **Control individual 1** | **Control individual 2** |
|---|---|---|---|---|---|
| Male/ female | F | F | F | F | M |
| Age (y) | 75 | 50 | 66 | 84 | 101 |
| Disease duration (y) | 4 | 5 | 3 | NA | NA |
| Clinical diagnosis | bvFTD | bvFTD | bvFTD | Control | Control |
| TDP-43 pathology | FTLD-TDP Type A | FTLD-TDP Type A | FTLD-TDP Type A | None | None |
| Tau pathology (Braak stage) | 0 | 0 | 1 | 0 | 1 |
| Amyloid-β pathology (Thal phase) | 1 | 0 | 3 | 0 | 0 |
| α-synuclein pathology | None | None | None | None | None |
| Disease-linked genetic variation | *GRN* c.1420_1421del | *GRN* c.1477C>T | *GRN* c.1414-2A>G | None | None |
| Reference | Case 5 in (Arseni *et al.* 2023 Nature 620, 898-903) | Case 2 in (Arseni *et al.* 2023 Nature 620, 898-903) (Arseni 2023) | Case 1 in (Cracco *et al.* 2022 Neuropath. Appl. Neurobiol. 48, e12836) | NA | NA |

F, female; M, male; y, years; bvFTD, behavioural-variant frontotemporal dementia; FTLD-TDP, frontotemporal lobar degeneration with TDP-43 pathology; NA, not applicable.
