## Supplementary Information for "Pathological TDP-43 filaments accumulate at synapses and cause synaptic dysfunction"

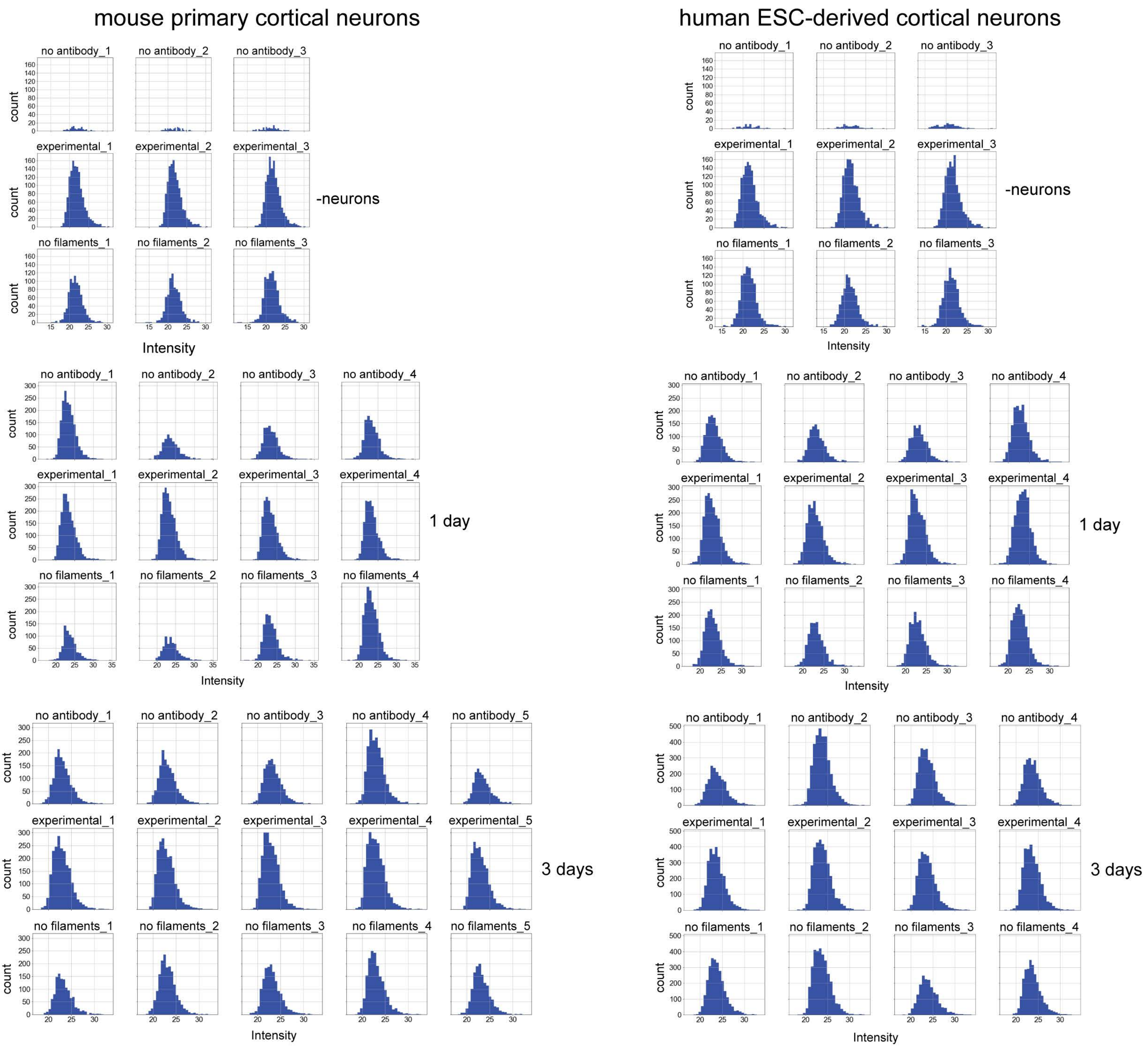


**Supplementary Information 1: Histograms of MS1 integrated signal intensities.** Counts of log2-transformed MS1 integrated signal intensities accross 25 bins for TDP-43 fiaments in the absence of neurons (-neurons), and for mouse primary neurons and human ESC-derived neurons incubated with TDP-43 filaments for 1 and 3 days. The first row shows the negative control group lacking the antibody (no antibody); the second row shows the experimental group (experimental); and the third row shows the negative control group lacking incubation with TDP-43 filaments (no filaments). Each column represents one biological replicate.

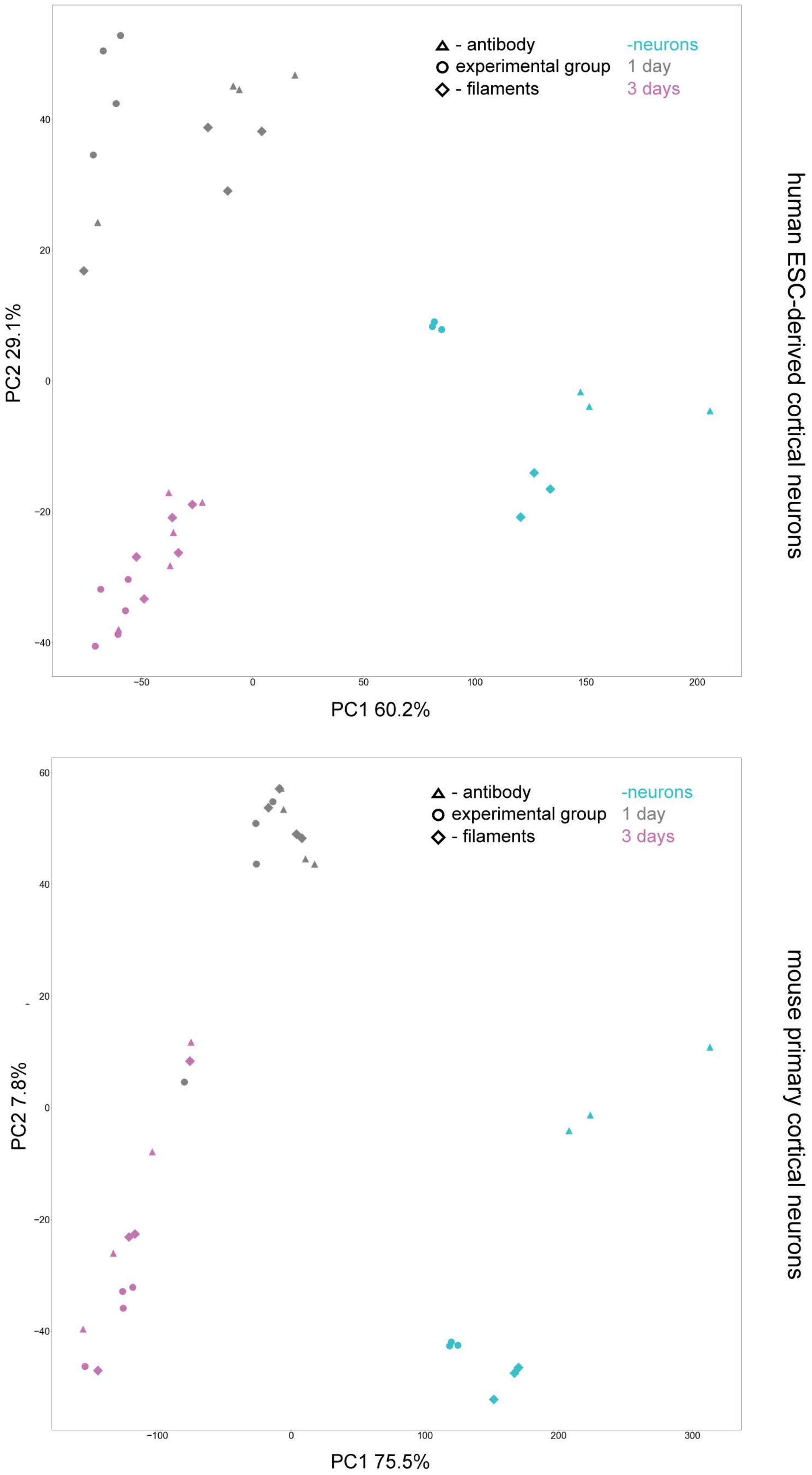


**Supplementary Information 2: Principal component analysis of imputated MS1 integrated signal intensities.** Plots of the first two principal components (PC1 and PC2) for imputated MS1 integrated signal intensities for the mouse primary cortical neurons and human ESC-derived cortical neurons. TDP-43 filaments in the absence of neurons are shown in cyan. Neurons incubated with TDP-43 filaments for 1 and 3 days are shown in grey and magenta, respectively. The negative control groups lacking the antibody (no antibody) and lacking the incubation with TDP-43 filaments (no filaments) are marked as triangles and diamonds, respectively. The experimental group (experimental) is marked as circles.

### Proximity labelling analysis pipeline

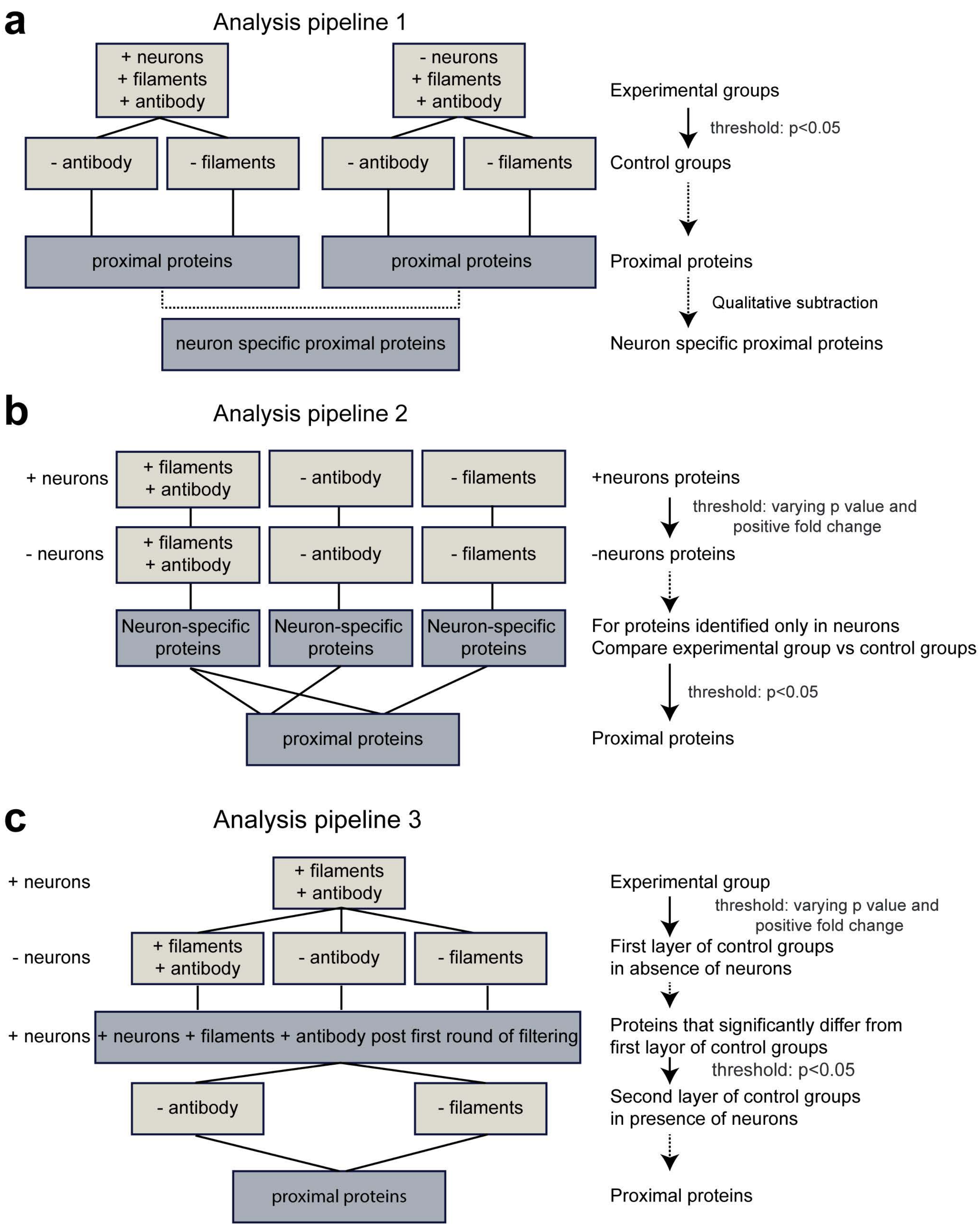


**Supplementary Information 3: Schematic of the three mass spectrometry data analysis pipelines. a,** In analysis pipeline 1, controlled proximity labelling experiments performed in the presence and absence of neurons were initially considered separately, followed by a qualitative comparison of the resulting proximal proteins to identify those specific to neurons. The control groups lacked antibody targeting (no antibody) and exogenous TDP-43 filaments (no filaments). **b,** In analysis pipeline 2, proteins from experimental, no antibody and no filaments controls in neurons were firstly filtered by using the equivalent experiments in the absence of neurons. The filtered experimental and control groups were then compared to identify proximal proteins. **c,** In analysis pipeline 3, proteins from the experimental group in the presence of neurons were filtered using the experimental and control groups in the absence of neurons. The resulting proteins were then compared to the control groups in the presence of neurons to identify proximal proteins.

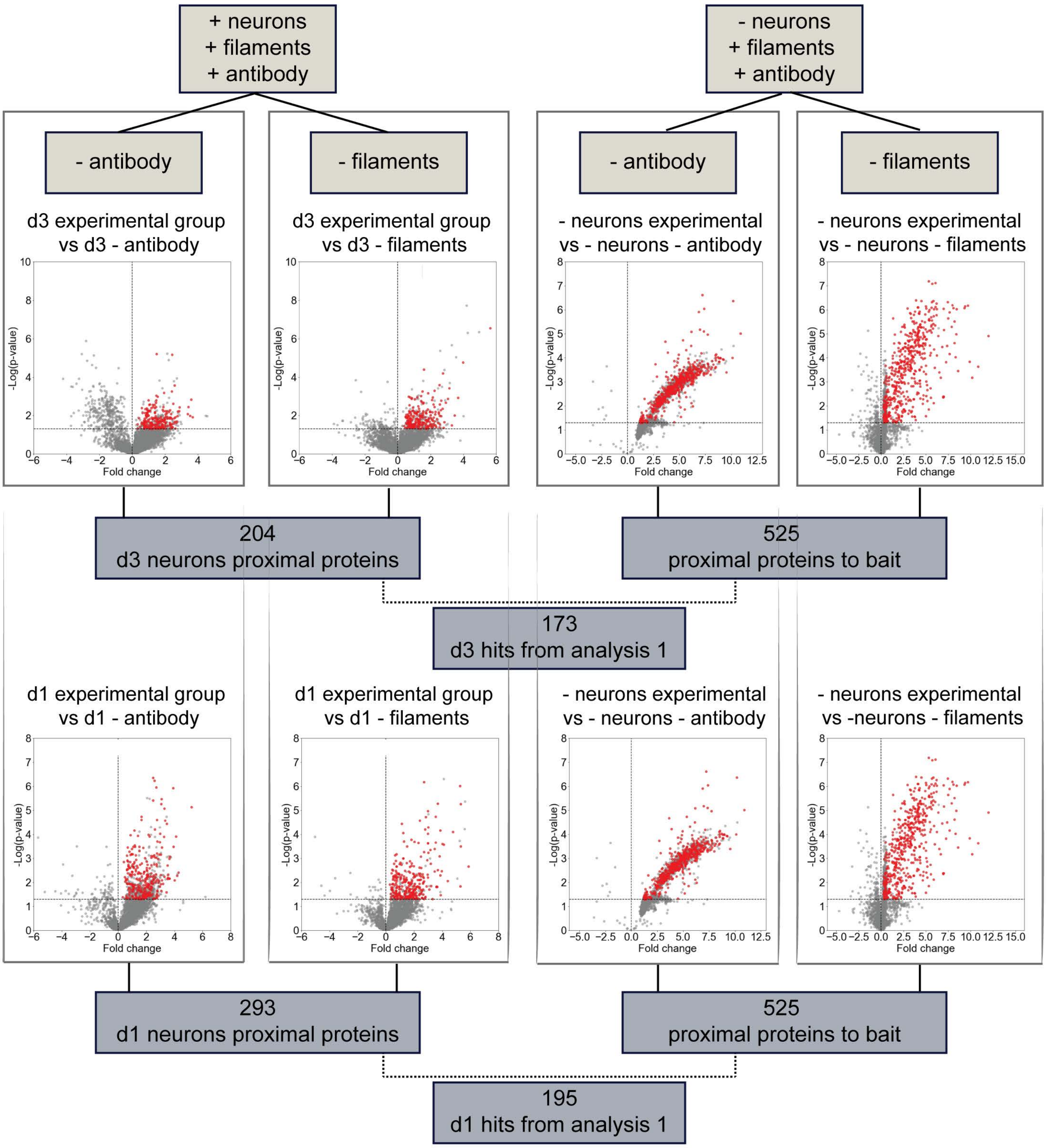


**Supplementary Information 4: Volcano plots of mass spectrometry data analysis pipeline 1 for mouse primary neurons.** Volcano plots comparing the quantitative MS data from experimental and control groups for proximity labelling carried out in mouse primary cortical neurons incubated with TDP-43 filaments for 1 d (bottom) and 3 d (top) using the first analysis pipeline. The fold change is plotted on the x-axis and the significance (-log10 transformed p value) is plotted on the y-axis. The dashed line marks the p value cut-off of 0.05. The selected proximal proteins (red) have p-values <0.05 and positive fold changes.

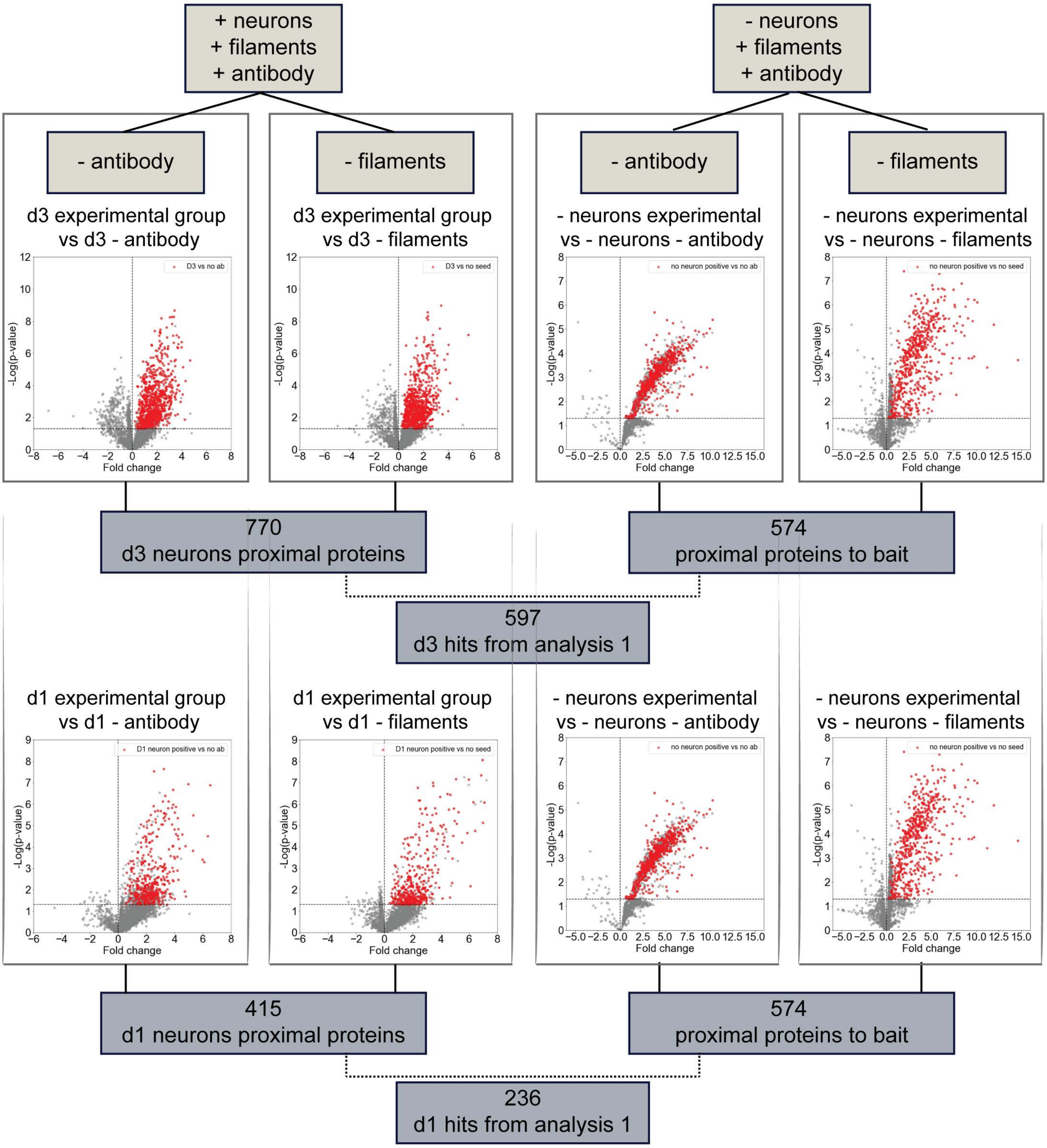


**Supplementary Information 5: Volcano plots of mass spectrometry data analysis pipeline 1 for human ESC-derived cortical neurons.** Volcano plots comparing the quantitative MS data from experimental and control groups for proximity labelling carried out in human ESC-derived cortical neurons incubated with TDP-43 filaments for 1 d (bottom) and 3 d (top) using the first analysis pipeline. The fold change is plotted on the x-axis and the significance (-log10 transformed p value) is plotted on the y-axis. The dashed line marks the p value cut-off of 0.05. The selected proximal proteins (red) have p-values <0.05 and positive fold changes.

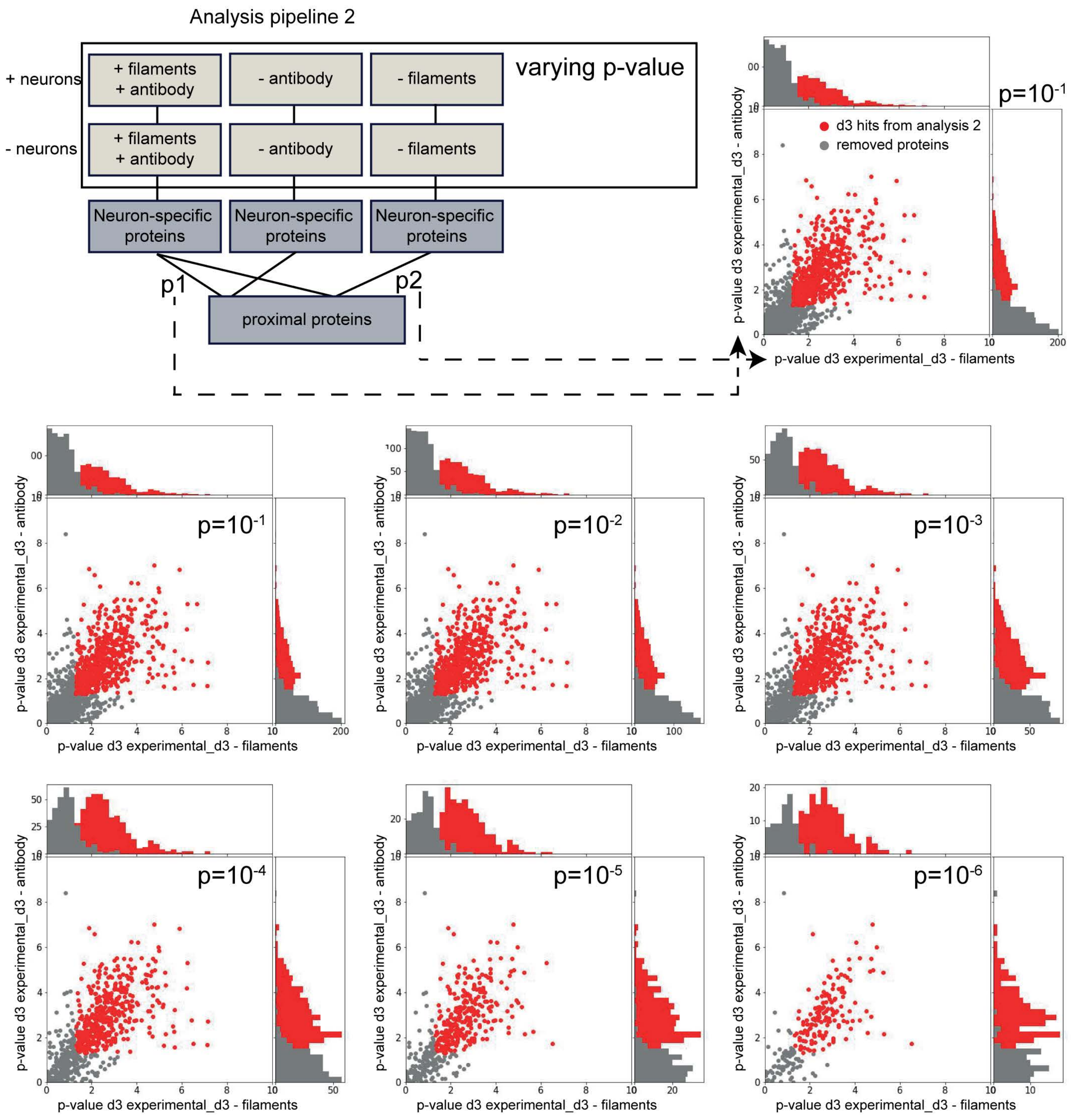


**Supplementary Information 6: Effect of p-value threshold in the filtering step of analysis pipeline 2.** The effect of varying the p-value threshold in the initial filtering step of analysis pipeline 2 was analysed by plotting p-values from comparing the filtered experimental group to filtered neuronal control groups lacking the antibody (P1) and TDP-43 filaments (P2). Histograms of the p-values within a set bin size for each comparison are shown. The p-value threshold used in the initial filtering step is shown in the top right corner of each scatter histogram plot. The selected proteins (red) have p-values $<0.05$. The analysis of human ESC-derived cortical neurons incubated with TDP-43 filaments for 3 days is shown.

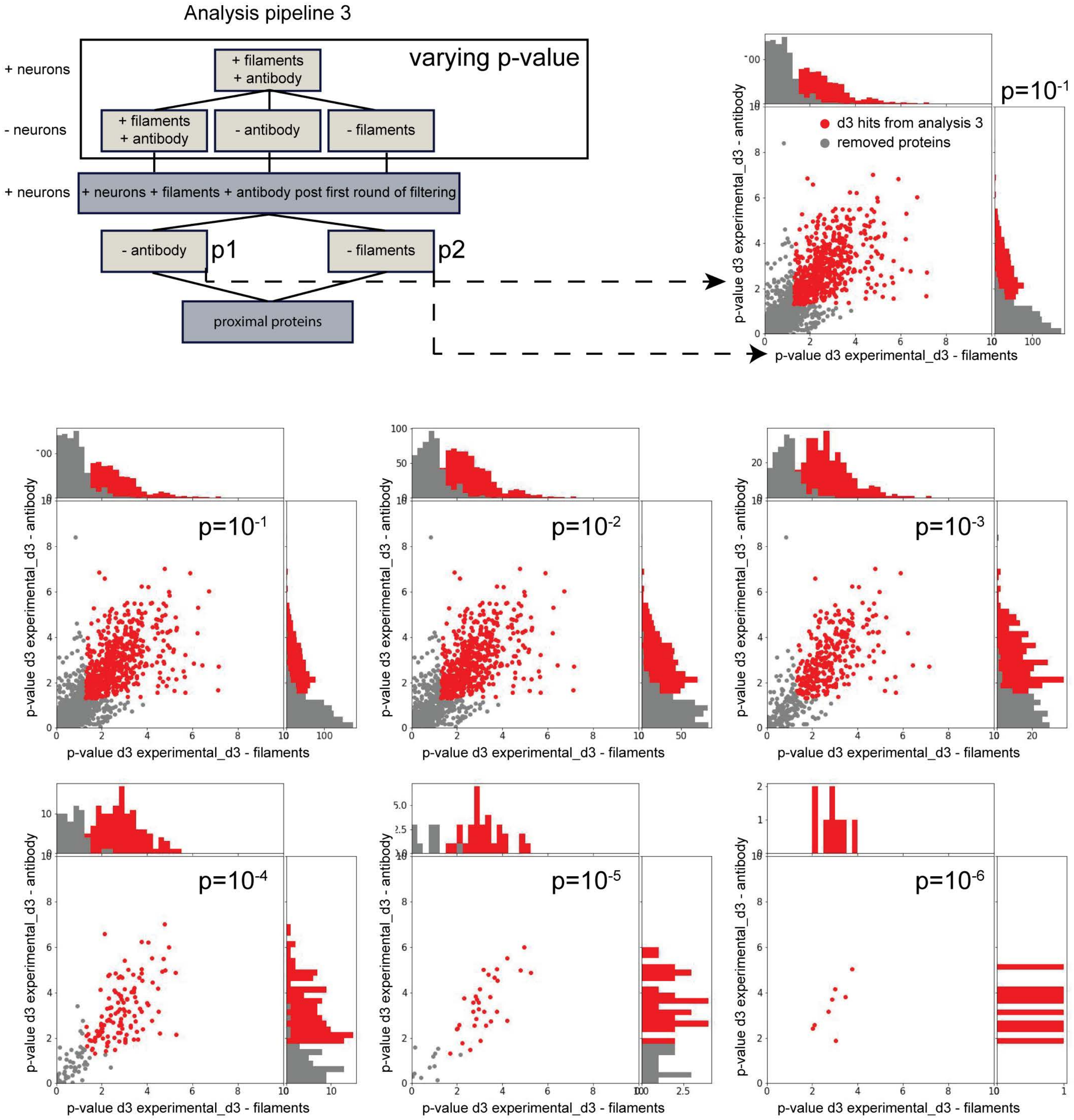


**Supplementary Information 7: Effect of p-value threshold in the filtering step of analysis pipeline 3.** The effect of varying the p-value threshold in the initial filtering step of analysis pipeline 3 was analysed by plotting p-values from comparing the filtered experimental group to neuronal control groups lacking antibody targeting (P1) and exogenous TDP-43 filaments (P2). Histograms of the p-values within a set bin size for each comparison are shown. The p-value threshold used in the initial filtering step is shown in the top right corner of each scatter histogram plot. The selected proteins (red) have p-values <0.05. The analysis of human ESC-derived cortical neurons incubated with TDP-43 filaments for 3 days is shown.

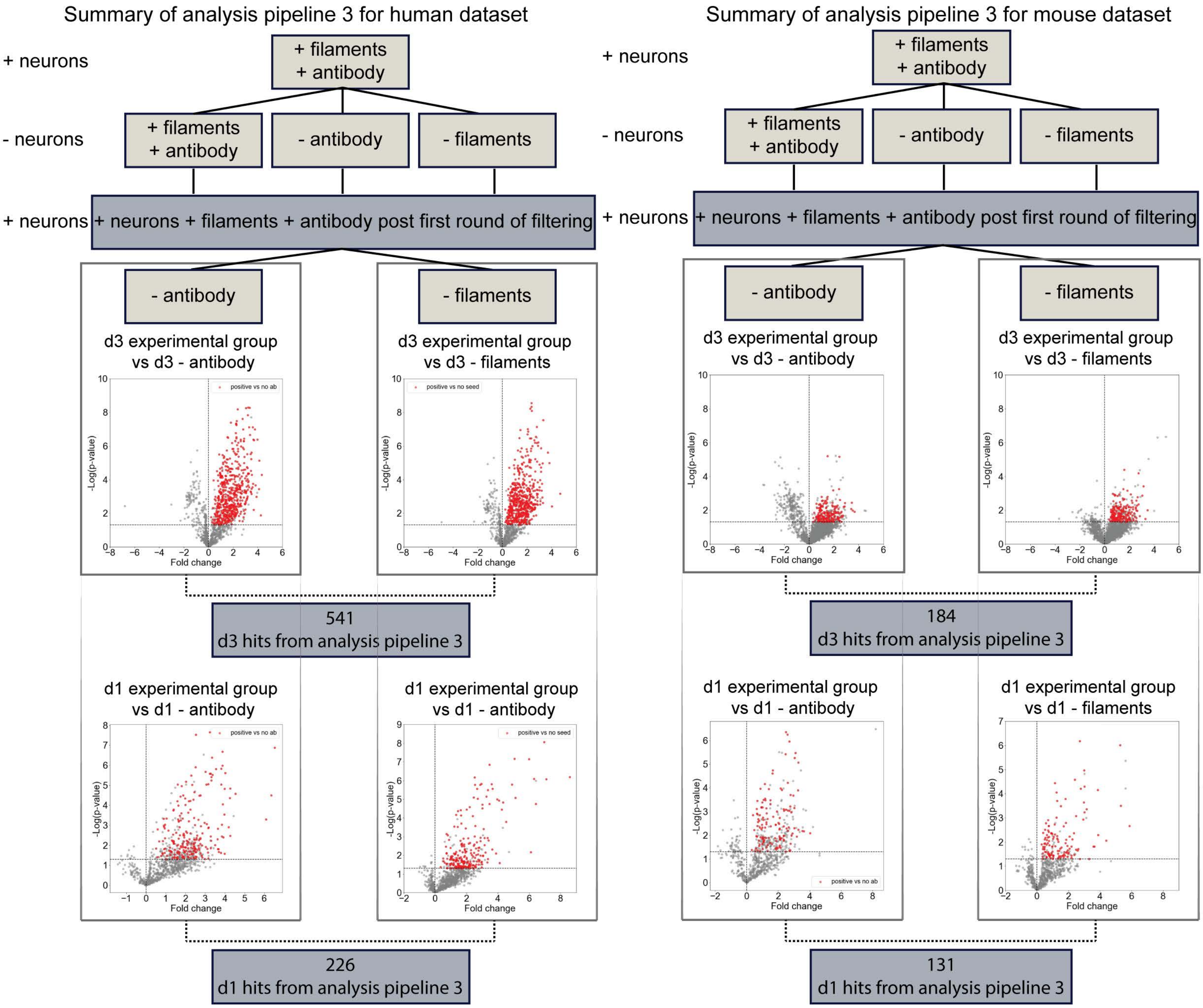


**Supplementary Information 8: Volcano plots of mass spectrometry data analysis pipeline 3.** Volcano plots comparing the quantitative MS data from experimental and control groups for proximity labelling carried out in mouse primary cortical neurons and human ESC-derived cortical neurons incubated with TDP-43 filaments for 1 and 3 days using the third analysis pipeline. The fold change is plotted on the x-axis and the significance (-log10 transformed p value) is plotted on the y-axis. The dashed line marks the p value cut-off of 0.05. The selected proximal proteins (red) have p-values <0.05 and positive fold changes.
